# JIP4 deficiency causes a lysosomal storage disease arising from impaired cystine efflux

**DOI:** 10.1101/2025.06.06.657909

**Authors:** Layla M. Nassar, Xiaojian Shi, Agnes Roczniak-Ferguson, Hongying Shen, Shawn M. Ferguson

**Affiliations:** Department of Cell Biology, Yale University School of Medicine, New Haven, Connecticut 06510, USA; Department of Neuroscience, Yale University School of Medicine, New Haven, Connecticut 06510, USA; Program in Cellular Neuroscience, Neurodegeneration and Repair, Yale University School of Medicine, New Haven, Connecticut 06510, USA; Wu Tsai Institute, Yale University School of Medicine, New Haven, Connecticut 06510, USA; Kavli Institute for Neuroscience, Yale University School of Medicine, New Haven, Connecticut 06510, USA; Department of Cellular and Molecular Physiology, Yale University School of Medicine, New Haven, Connecticut 06510, USA; Systems Biology Institute, Yale West Campus, West Haven, CT 06516, USA; Aligning Science Across Parkinson’s (ASAP) Collaborative Research Network, Chevy Chase, MD, 20815, USA

## Abstract

Lysosomes break down macromolecules, clear cellular waste and recycle nutrients such as cystine. We describe a novel mechanism whereby JIP4 regulates lysosomal cystine storage by controlling the abundance of cystinosin (CTNS), the transporter responsible for lysosomal cystine efflux. To this end, JIP4, previously characterized as a motor adaptor and kinase signaling scaffold, suppresses TMEM55B-dependent ubiquitylation of CTNS. Loss of JIP4 reduces CTNS protein levels, leading to lysosomal cystine accumulation and lysosomal storage defects that phenocopy loss of CTNS in both human cells and the renal proximal tubules of JIP4 knockout mice. These phenotypes mirror cystinosis, the lysosomal storage disease caused by CTNS loss-of-function. Our findings thus reveal a fundamental process that controls the efflux of lysosomal cystine and has relevance to understanding human disease arising from JIP4 mutations.

## Introduction

Lysosomes clear cellular waste by breaking down macromolecules and recycling nutrients (Ballabio & Bonifacino, 2020; Lawrence & Zoncu, 2019). When this degradative process or the subsequent efflux of metabolites is impaired, storage materials accumulate within lysosomes while the rest of the cell experiences nutrient deprivation (Laqtom et al., 2022). The physiological burden of aberrant metabolite recycling from lysosomes, which is exemplified by at least 70 characterized lysosomal storage diseases (Platt et al., 2018), illustrates the importance of cellular mechanisms that coordinate lysosome degradative activity with nutrient recycling.

Lysosomes face enormous challenges in managing levels of the conditionally essential amino acid cysteine. Proteolysis within lysosomes liberates cysteine, which is subsequently oxidized to form cystine before being transported to the cytoplasm where it is reduced back into cysteine (Gahl, Bashan, et al., 1982; Pisoni et al., 1990). In addition to supporting protein translation, cytoplasmic cysteine is the rate-limiting component of glutathione, a major antioxidant, and therefore critical to the cellular response to oxidative stress and reversal of lipid peroxidation that leads to ferroptosis (Rathbun & Murray, 1991; Stipanuk et al., 2006; Jiang et al., 2021; Poltorack & Dixon, 2022)). Simultaneously, overload of cytoplasmic cysteine results in toxicity by altering iron homeostasis, requiring the sequestration of cysteine within the lysosome (Hughes et al., 2020). Lysosomal cysteine is also required for thiol reduction of disulfide bonds as well as the activity of cysteine cathepsin proteases (Mego, 1984; Lloyd, 1986; Arunachalam et al., 2000). Therefore, proteins cannot be efficiently degraded within lysosomes without correctly managing levels of reduced lysosomal cysteine. The need to tightly control the compartmentalization of cysteine between lysosomes and the cytoplasm may explain why, in addition to the cysteine liberated by proteolysis within lysosomes, cysteine is additionally transported into lysosomes by Mfsd12 (Adelmann et al., 2020). Meanwhile, the efflux of cystine from lysosomes is mediated by cystinosin (CTNS) (Kalatzis et al., 2001).

Loss-of-function mutations in CTNS cause a lysosomal storage disease known as cystinosis (Town et al., 1998; Gahl et al., 2002; Jamalpoor et al., 2021). Cystinosis is characterized by the accumulation of cystine within lysosomes and widespread organ dysfunction. In nephropathic cystinosis, the kidney is particularly affected, as CTNS deficiency induces proximal tubule dysfunction (Fanconi syndrome) and eventual end-stage renal failure, leading most cystinosis patients to eventually require renal transplantation for survival (Elmonem et al., 2016). Accumulation of cystine impairs both normal lysosome functions and limits cellular cysteine availability (Gahl, Bashan, et al., 1982; Cherqui et al., 2002; Kalatzis et al., 2004; Jouandin et al., 2022; Berquez et al., 2023). CTNS-deficient cells upregulate transcription factors involved in antioxidant response, accumulate lipid peroxides, and become hypersensitive to ferroptotic cell death, pointing to a role in maintaining antioxidant capacity (Armenta et al., 2022; He et al., 2023; Swanda et al., 2023; Yu et al., 2024). Collectively, these findings emphasize the importance of lysosomal cystine sequestration and release in supporting cellular cysteine and redox balance. However, little is known about the regulatory mechanisms that tune CTNS activity to dynamic cellular demands.

Here, we reveal a novel mechanism for regulating lysosomal cystine efflux through JNK-interacting protein 4 (JIP4, encoded by the SPAG9 gene), a protein that was originally identified as a scaffold for components of the p38 MAPK signaling pathway and which is best characterized as linking lysosomes to molecular motors (Kelkar et al., 2005; Montagnac et al., 2009; Willett et al., 2017; Gowrishankar et al., 2021; Celestino et al., 2022; Cason & Holzbaur, 2023). We now show that JIP4 ensures lysosomal cystine export by counteracting the ubiquitin-dependent degradation of CTNS. We identify TMEM55B as an adaptor for CTNS ubiquitylation that is negatively regulated by JIP4. We further demonstrate that, in the absence of JIP4, increased ubiquitylation accelerates CTNS degradation and results in *in vivo* phenotypes that resemble cystinosis. Observations from human cell culture to mouse models thus establish that JIP4 coordinates lysosomal abundance of CTNS with cellular metabolic demands. These findings broaden understanding of lysosomal function, define a new pathway that results in lysosomal storage disease and shed light on how aberrant cystine storage may contribute to human disease arising from JIP4 loss-of-function mutations.

## Results

Extending from previous observations of lysosome mislocalization following siRNA-mediated depletion of JIP4, we saw a striking accumulation of lysosomes at the periphery of a newly generated line of JIP4 KO HeLa cells (Extended Data Fig. 1; Willett et al., 2017). Unexpectedly, over the course of 2-3 days following each passaging of the JIP4 KO cells, vacuoles appeared that were readily visualized by brightfield microscopy as the most prominent feature in these cells (Fig. 1a,b). These vacuoles contained the integral membrane protein LAMP1 and therefore represent enlarged late endosomes or lysosomes (Fig. 1c, d). To test whether accumulation of lysosomes in peripheral cellular protrusions was sufficient to induce this massive lysosome enlargement, we over-expressed SKIP, a scaffold that links lysosomes to kinesin and promotes their accumulation at the cell periphery (Rosa-Ferreira & Munro, 2011). Although SKIP over-expression caused lysosomes to accumulate at the cell periphery in a similar manner to what occurred in JIP4 KO cells, this redistribution did not lead to their enlargement (Extended Data Fig. 2).

**Fig. 1:**
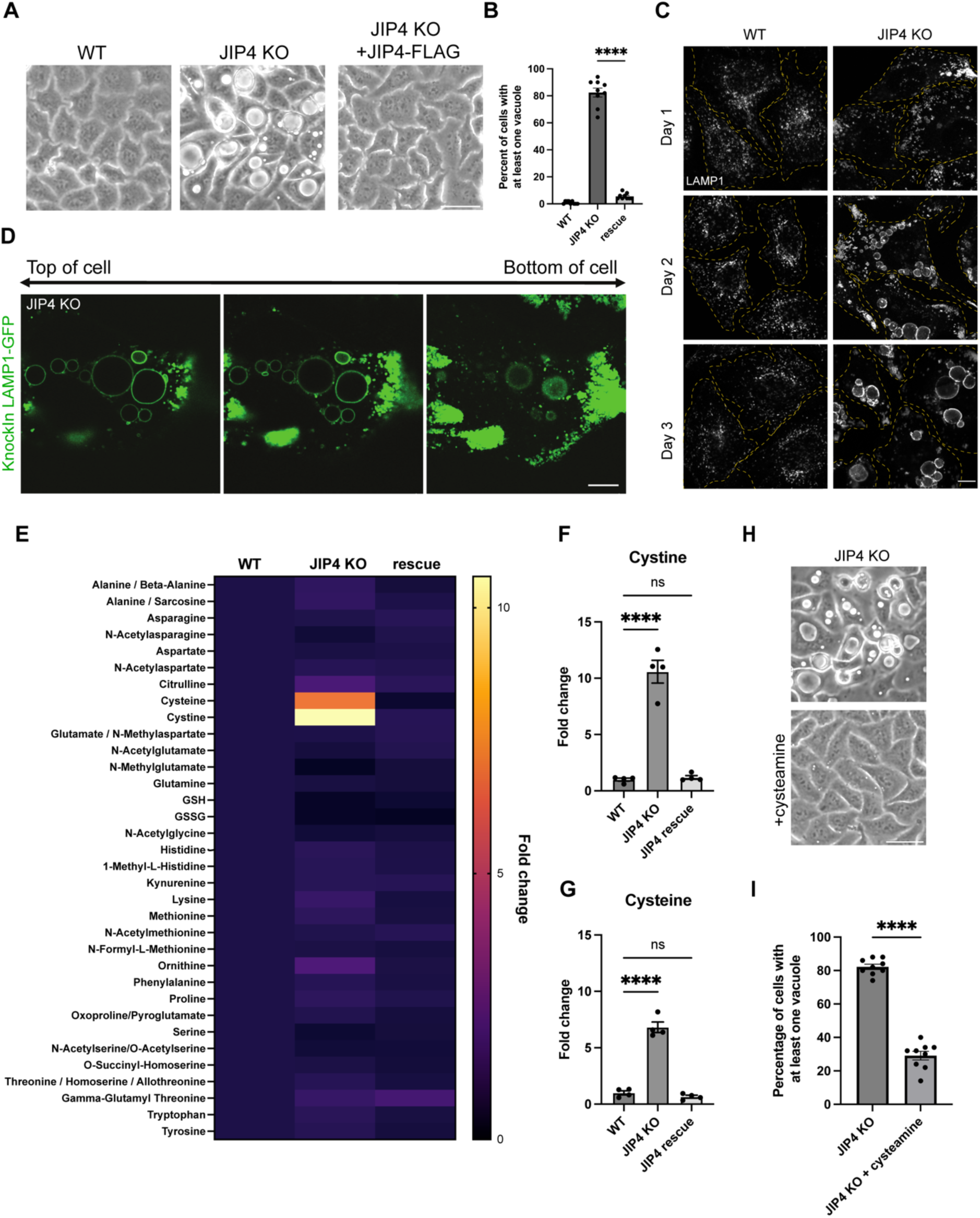
JIP4 KO cells aberrantly accumulate cysteine and cystine. **a,** Brightfield imaging of wildtype (WT), JIP4 KO, and rescue (JIP4 KO +JIP4-FLAG) HeLa cells three days post-plating. Scale bar = 50 µm. **b,** Quantification of WT, JIP4 KO, and JIP4 KO +JIP4-FLAG cells in **a** with at least one vacuole visible by brightfield (mean ± SEM, n=3 independent experiments, ****P<0.0001, one-way ANOVA with Tukey’s multiple comparisons test). **c,** Immunofluorescent staining for LAMP1 at day 1, 2, and 3 post-plating in WT and JIP4 KO cells. Scale bar = 10 µm. **d,** Images of JIP4 KO cells with LAMP1-GFP endogenous tag at 3 different z-planes. Scale bar = 10 µm. **e,** Results from HILIC-amide mass spectrometry of cell pellets from WT, JIP4 KO, and JIP4 KO +JIP4-FLAG cells. Fold change average calculated from four biological replicates. **f-g,** Fold change in cystine **(f)** and cysteine **(g)** levels (n=4 biological replicates, ****P<0.0001, one-way ANOVA with Tukey’s multiple comparisons test). **h**, Brightfield imaging of JIP4 KO cells with and without the addition of 1mM cysteamine. Scale bar = 50 µm. **i,** Quantification of JIP4 KO cells in **h** with at least one vacuole visible by brightfield (mean ± SEM, n=3 independent experiments, ****P<0.0001, unpaired *t* test)

Interestingly, changing cell culture media daily prevented vacuoles from forming in JIP4 KO cells, indicating that nutrient depletion might be a driver of their formation (Extended Data Fig. 3). Additionally, the fact that vacuoles still formed when media changes were supplemented with dialyzed FBS, which removes components under 10 kDa, instead of complete FBS suggested that a low molecular weight metabolite was responsible for preventing vacuole formation (Extended Data Fig. 3). Because a connection between JIP4 and metabolism had not been previously identified, we screened for changes in common amino acids by performing mass-spec based polar metabolic profiling in wild type (WT), JIP4 KO, and JIP4 KO + JIP4-FLAG (JIP4 rescue) cells. Cystine, the oxidized form of the amino acid cysteine that is predominantly found in lysosomes, accumulated in JIP4 KO cells by over 10-fold, and cysteine accumulated by over 5-fold (Fig. 1e-g; Schulman et al., 1969; Gahl, Bashan, et al., 1982; Abu-Remaileh et al., 2017). However, no other metabolites showed comparable changes. These results suggested that JIP4 KO cells have a defect in the lysosomal efflux of cystine. To test whether increasing lysosomal cystine efflux would alleviate vacuole formation, we added the drug cysteamine which is used clinically for the treatment of cystinosis. Cysteamine facilitates lysosomal cystine export by forming cysteine-cysteamine mixed disulfides that resemble lysine and thus exit the lysosome via PQLC2, the lysosomal transporter for cationic amino acids (Fig. 1h,I; Gahl et al., 1985; Jezegou et al., 2012). Cysteamine treatment reduced the abundance of vacuoles in JIP4 KO cells, suggesting that vacuoles arise due to aberrant lysosomal cystine sequestration. This unexpected connection between JIP4 and cystine metabolism raised questions on how JIP4 intersects with cystine metabolism.

We next knocked out the lysosomal cystine exporter cystinosin (CTNS), as CTNS--depleted cells are known to massively accumulate cystine in a similar manner to what we observed in JIP4 KO (Gahl, Bashan, et al., 1982; Gahl, Tietze, et al., 1982; Park et al., 2002). CTNS KO was also previously reported to cause lysosome enlargement in cultured cells (Hollywood et al., 2020). This led to the observation that CTNS KO HeLa cells were just like JIP4 KO cells with respect to the formation of massively enlarged lysosomes 2-3 days following each passaging (Fig. 2a,b). To further investigate the relationship between lysosome cystine accumulation and lysosome enlargement in CTNS KO and JIP4 KO cells, we again treated cells with cysteamine (Gahl et al., 1985; Jezegou et al., 2012). Cysteamine treatment suppressed the enlarged lysosome phenotype in both JIP4 KO and CTNS KO cells, suggesting that enlarged lysosomes arise due to defects in CTNS-mediated cystine efflux (Fig. 2c,d). Consistent with a defect in CTNS-mediated lysosome cystine efflux being a major consequence of JIP4 depletion, overexpression of CTNS-FLAG rescued the striking lysosome enlargement in JIP4 KO cells (Fig. 2e,f).

**Fig. 2:**
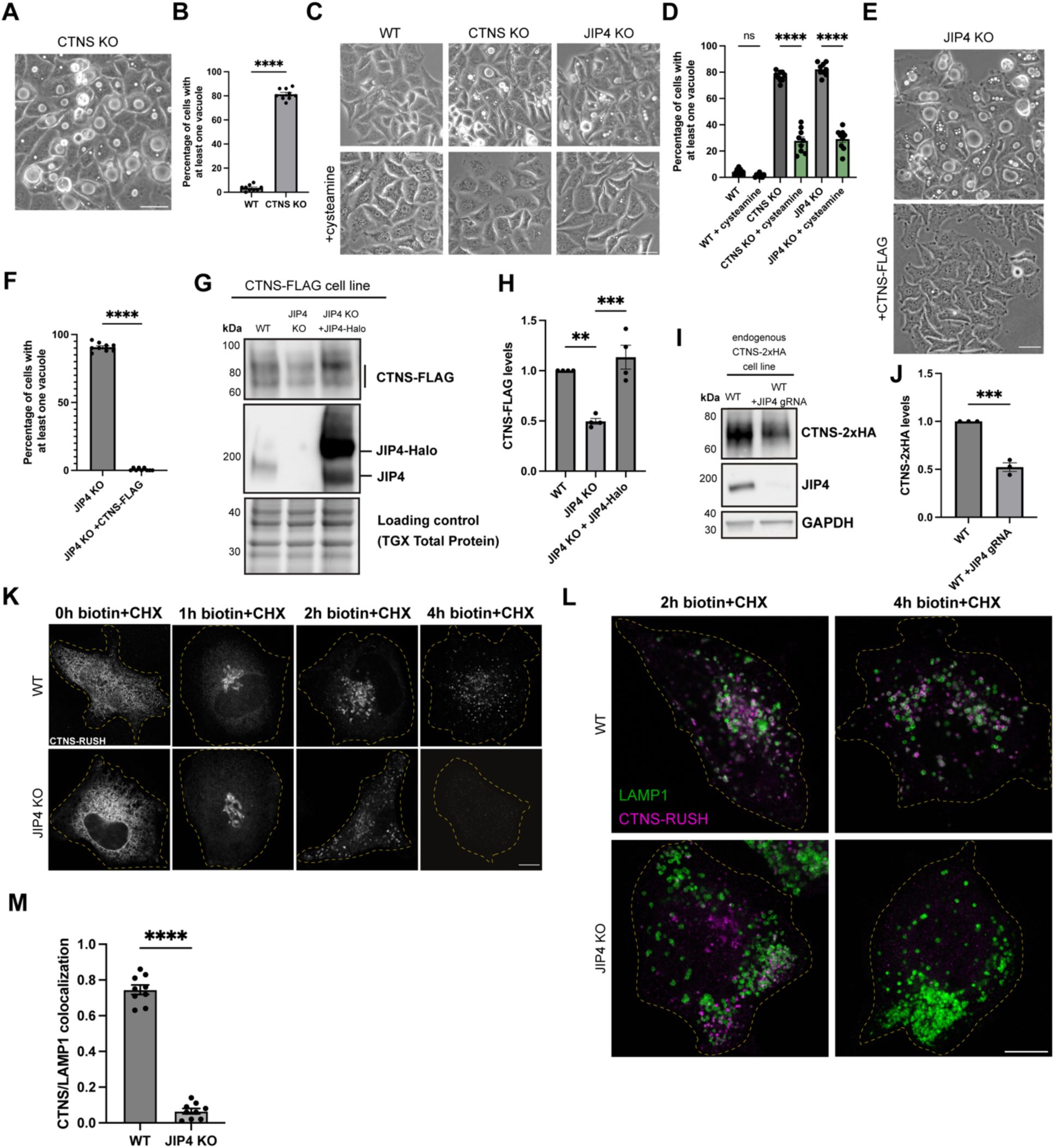
JIP4 stabilizes cystinosin (CTNS) at lysosomes. **a,** Brightfield imaging of CTNS KO cells after 3 days post-plating. Scale bar = 50 µm. **b,** Quantification of the percentage of WT and CTNS KO cells with at least one vacuole visible by brightfield (mean ± SEM, n=3 independent experiments, ****P<0.0001, unpaired *t* test). **c,** Brightfield imaging of WT, JIP4 KO, and CTNS KO cells with and without 1mM cysteamine three days post-plating. Scale bar = 50 µm. **d,** Quantification of the percentage of WT, JIP4 KO, and CTNS KO cells in **c** with at least one vacuole visible by brightfield (mean ± SEM, n=3 independent experiments, ****P<0.0001, one-way ANOVA with Tukey’s multiple comparisons test). **e,** Brightfield imaging of JIP4 KO cells with and without stable overexpression of CTNS-FLAG. Scale bar = 50 µm. **f,** Quantification of JIP4 KO and JIP4 KO +CTNS-FLAG cells in **e**, n=3 biological replicates (mean ± SEM, n=3 independent experiments, ****P<0.0001, unpaired *t* test). **g,** Western blot analysis of WT +CTNS-FLAG, JIP4 KO +CTNS-FLAG, and JIP4 KO +CTNS-FLAG with transient transfection of JIP4-Halo. **h,** Quantification of CTNS-FLAG levels in **g** (mean ± SEM, n=4 independent experiments, ***P=0.0003, **P=0.0017, one-way ANOVA with Tukey’s multiple comparisons test). **i**, Western blot analysis of HeLa cells with endogenously tagged CTNS-2xHA, with and without transfection of JIP4 gRNAs. **j,** Quantification of CTNS-2xHA levels in **i** (mean ± SEM, n=3 independent experiments, ***P=0.0004, unpaired *t* test). **k,** CTNS-RUSH construct in WT and JIP4 KO cells with biotin + cycloheximide (CHX) at indicated times given. Scale bar = 10 µm. **l,** CTNS-RUSH and LAMP1 in WT and JIP4 KO cells after 2h biotin+CHX and 4h biotin+CHX. Scale bar = 10 µm. **m,** Quantification of CTNS-LAMP1 colocalization after 4h of biotin+CHX treatment (mean ± SEM, n=3 independent experiments, ****P<0.0001, unpaired *t* test).

To further investigate mechanisms linking JIP4 and CTNS, JIP4 was depleted from HeLa cells that stably expressed FLAG-tagged CTNS (CTNS-FLAG). JIP4 KO cells had significantly lower levels of CTNS-FLAG protein, and the reduction in CTNS-FLAG was rescued upon introduction of Halo-tagged JIP4 (Fig. 2g,h). This impact of JIP4 depletion on CTNS protein levels was independently observed in knockin cells with 2xHA epitope-tagged endogenous CTNS protein (Fig. 2i,j). To better understand the mechanism underlying JIP4-dependent control of CTNS protein levels, we used the Retention Using Selective Hooks (RUSH) strategy to observe the trafficking of newly made CTNS to lysosomes (Boncompain et al., 2012). The RUSH system allows for endoplasmic reticulum retention of tagged CTNS until biotin is added. We additionally co-applied cycloheximide (CHX) to block new protein synthesis. This pulse-chase strategy revealed that CTNS-RUSH initially trafficked similarly in both WT and JIP4 KO cells as it sequentially exhibited an ER-like distribution before biotin+CHX, a Golgi-like distribution after 1h with biotin+CHX, and a predominantly endosome-like punctate distribution after 2h biotin+CHX. However, after 4h, CTNS was abundant on lysosomes in WT cells but difficult to detect in JIP4 KO cells (Fig. 2k-m). This result indicated that, in the absence of JIP4, CTNS is rapidly degraded upon arrival at lysosomes.

In budding yeast, control of vacuolar transporter stability occurs via ubiquitylation and direct internalization into the vacuole/lysosome in a process termed ESCRT-mediated degradation (Li et al., 2015; Zhu et al., 2017). This regulation applies to members of the PQ-loop family that are distantly related to CTNS. We therefore hypothesized that aberrant ubiquitylation was responsible for the degradation of CTNS in JIP4 KO lysosomes. To test this, the three cytoplasmic facing lysines in CTNS were mutated to alanine (K->A) to prevent ubiquitylation (Fig. 3a). Immunoprecipitation of CTNS-FLAG and the K->A mutant revealed that WT CTNS-FLAG is modified by HA-Ubiquitin, but the K->A mutant is not. Additionally, after normalizing for the reduced abundance of CTNS-FLAG in JIP4 KO cells, CTNS-FLAG was more ubiquitylated in the absence of JIP4 (Fig. 3b,c). The K->A mutant of CTNS-RUSH rescued CTNS localization at the lysosome following biotin treatment in JIP4 KO cells (Fig. 3d,e). Stable overexpression of CTNS-FLAG K->A in JIP4 KO cells also rescued vacuole formation (Fig. 2e, Extended Data Fig. 4). To test whether loss of CTNS was due to degradation at the lysosome, lysosomal acidification was inhibited with folimycin, an inhibitor of the V-ATPase. This treatment rescued CTNS levels in JIP4 KO cells (Fig. 3f,g). Together, these data are consistent with a JIP4-dependent inhibition of CTNS ubiquitylation which prevents CTNS degradation in lysosomes.

**Fig. 3:**
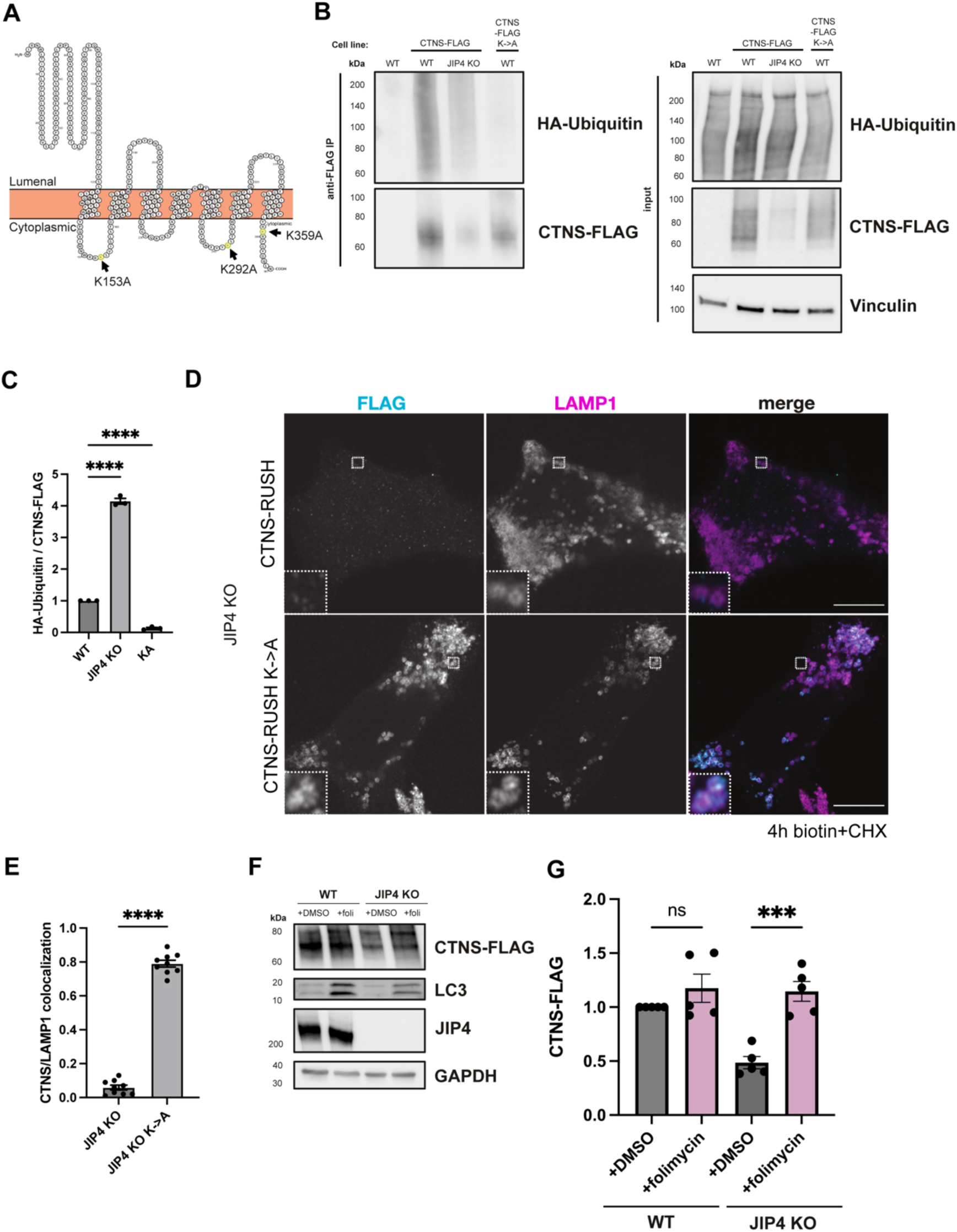
JIP4 suppresses ubiquitylation and degradation of CTNS. **a,** Protter diagram displaying the transmembrane topology of CTNS with its three cytoplasmic facing lysines (K) highlighted. **b,** Western blot analysis of anti-FLAG immunoprecipitation in WT, WT +CTNS-FLAG, JIP4 KO +CTNS-FLAG, and WT +CTNS-FLAG-K->A cells. **c,** Quantification of CTNS-FLAG levels in **b** (mean ± SEM, n=3 independent experiments, ****P<0.0001, one-way ANOVA with Tukey’s multiple comparisons test). **d,** CTNS-RUSH and CTNS-RUSH-K->A and LAMP1 JIP4 KO cells at 4h biotin+CHX. Scale bar = 10 µm. **e,** Quantification of CTNS/LAMP1 colocalization at 4h biotin+CHX in **k** (mean ± SEM, n=3 independent experiments, ****P<0.0001, unpaired *t* test). **f,** Western blot analysis of WT +CTNS-FLAG and JIP4 KO +CTNS-FLAG with and without folimycin. **g,** Quantification of CTNS-FLAG levels in **f** (mean ± SEM, n=3 independent experiments, *P=0.0191, **P=0.0035, one-way ANOVA with Tukey’s multiple comparisons test).

How does JIP4 prevent CTNS ubiquitylation? Recent work showed that a JIP4 interacting protein, TMEM55B, binds several members of the NEDD4-like family of E3 ubiquitin ligases through its PY-containing motif and could direct the ubiquitylation of substrates at lysosomes (Jeong et al., 2024). We hypothesized that TMEM55B may be acting similarly to the yeast vacuolar Ssh4, a PY motif-containing adaptor that facilitates the ubiquitylation of the yeast vacuolar exporter Ypq1 by Rsp5 (Li et al., 2015; Zhu et al., 2017; Arines et al., 2021); TMEM55A/B and Ssh4 share cytoplasmic facing PY motifs that can recruit NEDD4 ubiquitin ligases to the lysosomal/vacuolar surface (Extended Data Fig. 5a, b)(Jeong et al., 2024; Jumper et al., 2021; Varadi et al., 2023). We thus formed a model in which TMEM55B acts as an adaptor to facilitate CTNS ubiquitylation. Consistent with this hypothesis, knockdown of TMEM55B and its homolog TMEM55A increased levels of CTNS, while knockdown of TMEM55A/B in CTNS K->A mutant cells did not alter levels of CTNS, indicating a TMEM55A/B-dependent depletion that relies on ubiquitylation (Fig. 4a,b). Additionally, immunoprecipitation of CTNS-FLAG revealed copurification of TMEM55B but not the abundant lysosomal transmembrane protein LAMP1 (Fig. 4c). To test the role of JIP4 in this interaction, CTNS-FLAG was immunoprecipitated in both control and JIP4-Halo overexpressing cells, and we observed that JIP4 over-expression resulted in less CTNS interaction with TMEM55B (Fig. 4d,e). This observation suggested JIP4 suppresses TMEM55B-facilitated ubiquitylation of CTNS. To further test this model and to identify whether NEDD4-like E3 ligases contribute to this ubiquitylation, mCherry-TMEM55B was immunoprecipitated, and samples were blotted for NEDD4. Co-immunoprecipitation of NEDD4 with mCherry-TMEM55B significantly increased in JIP4 KO cells compared to WT cells, showing that JIP4 inhibits the NEDD4-TMEM55B interaction (Fig. 4f,g). Together, this supports a model in which JIP4 inhibits TMEM55B from promoting the ubiquitylation of CTNS by NEDD4 (and potentially additional close homologs to NEDD4), and CTNS’s subsequent depletion from the lysosome (Fig. 4h).

**Fig. 4:**
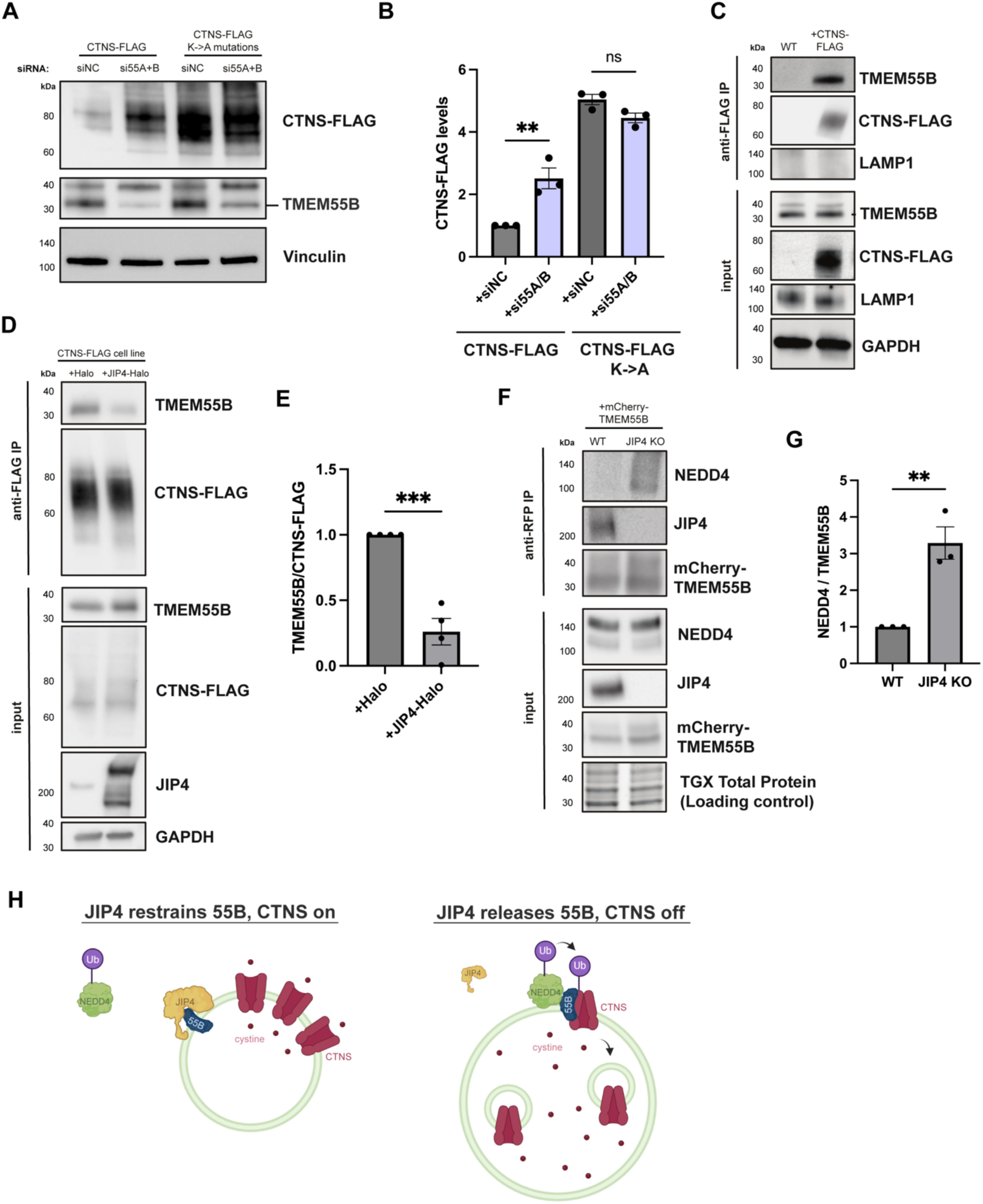
JIP4 protects CTNS from TMEM55B-mediated degradation. **a,** Western blot analysis of CTNS-FLAG and CTNS-FLAG-K->A with and without siRNA depletion of TMEM555A and TMEME55B. **b,** Quantification of CTNS-FLAG levels in **a** (mean ± SEM, n=5 independent experiments, *P=0.0191, **P=0.0038, one-way ANOVA with Tukey’s multiple comparisons test). **c,** Western blot analysis of anti-FLAG immunoprecipitation in WT and WT +CTNS-FLAG cells. **d,** Western blot analysis of anti-FLAG immunoprecipitation in WT +CTNS-FLAG cells with and without over-expression (transient transfection) of JIP4-Halo. **e,** Quantification of TMEM55B/CTNS-FLAG levels in **d**, (mean ± SEM, n=3 independent experiments, *P=0.0292, unpaired *t* test). **f,** Western blot analysis of anti-RFP immunoprecipitation in WT and JIP4 KO with transient transfection of mCherry-TMEM55B. **g,** Quantification of NEDD4/TMEM55B in WT and JIP4 KO cells in **f** (mean ± SEM, n=3 independent experiments, **P=0.0089, paired *t* test). **h,** Model for JIP4-mediated suppression of TMEM55B-dependent ubiquitylation of CTNS.

To establish a physiological understanding of JIP4-dependent regulation of CTNS, a new strain of JIP4 knockout mice was created via CRISPR-mediated deletion of exon 4 (Extended Fig. 6a-c). Strikingly, JIP4 knockout mice had lighter skin and hair color than their wildtype and heterozygous littermates (Extended Fig. 6d). This was similar to the impact of CTNS mutations, which cause an analogous hypopigmentation in both mice and humans (Chiaverini et al., 2012). This hypopigmentation arises due to the role played by cysteine in controlling the balance between synthesis of eumelanin and pheomelanin in melanosomes, a lysosome-related organelle that is responsible for pigment synthesis in melanocytes (Borovanský & Riley, 2011; Chiaverini et al., 2012).

Kidney failure arising from renal proximal tubule dysfunction represents the most severe phenotype arising in humans with loss-of-function mutations in CTNS. Likewise, CTNS knockout mice proximal tubules exhibit lysosome vacuolization and cystine crystal formation in mice aged 12 months and above (Cherqui et al., 2002). Therefore, defects within the renal proximal tubule were next investigated in 12-to 14-month-old WT and JIP4 KO mice. The renal proximal tubule undergoes extreme levels of endocytosis to remove proteins from urine, and lysosomes are enriched directly under the apical membrane microvilli to quickly capture and degrade endocytosed proteins (Fig. 5a; Rodman et al., 1986). JIP4 is present on these apically enriched lysosomes (Extended Data Fig. 7a). In contrast to the strong apical enrichment of lysosomes in WT kidneys, lysosomes were more abundant along the basolateral membrane in JIP4 KO proximal tubules, indicating that JIP4 is critical for the placement of lysosomes in the proximal tubule (Fig. 5b, e; Extended Data Fig. 7b). However, the endocytic markers clathrin light chain (CLC) and megalin shared similar apical enrichment in both WT and JIP4 KO cells (Extended Data Fig. 7c,d). These results demonstrate that while JIP4 is not required for development and maintenance of proximal tubule polarity, late endosomes and lysosomes are selectively dependent on JIP4 for their enrichment under the apical microvilli. This data is consistent with a role for JIP4 in minus end directed movement of lysosomes within proximal tubule cells, as minus ends of microtubules are polarized towards the apical membrane (Bacallao et al., 1989; Dammermann et al., 2003).

**Fig. 5:**
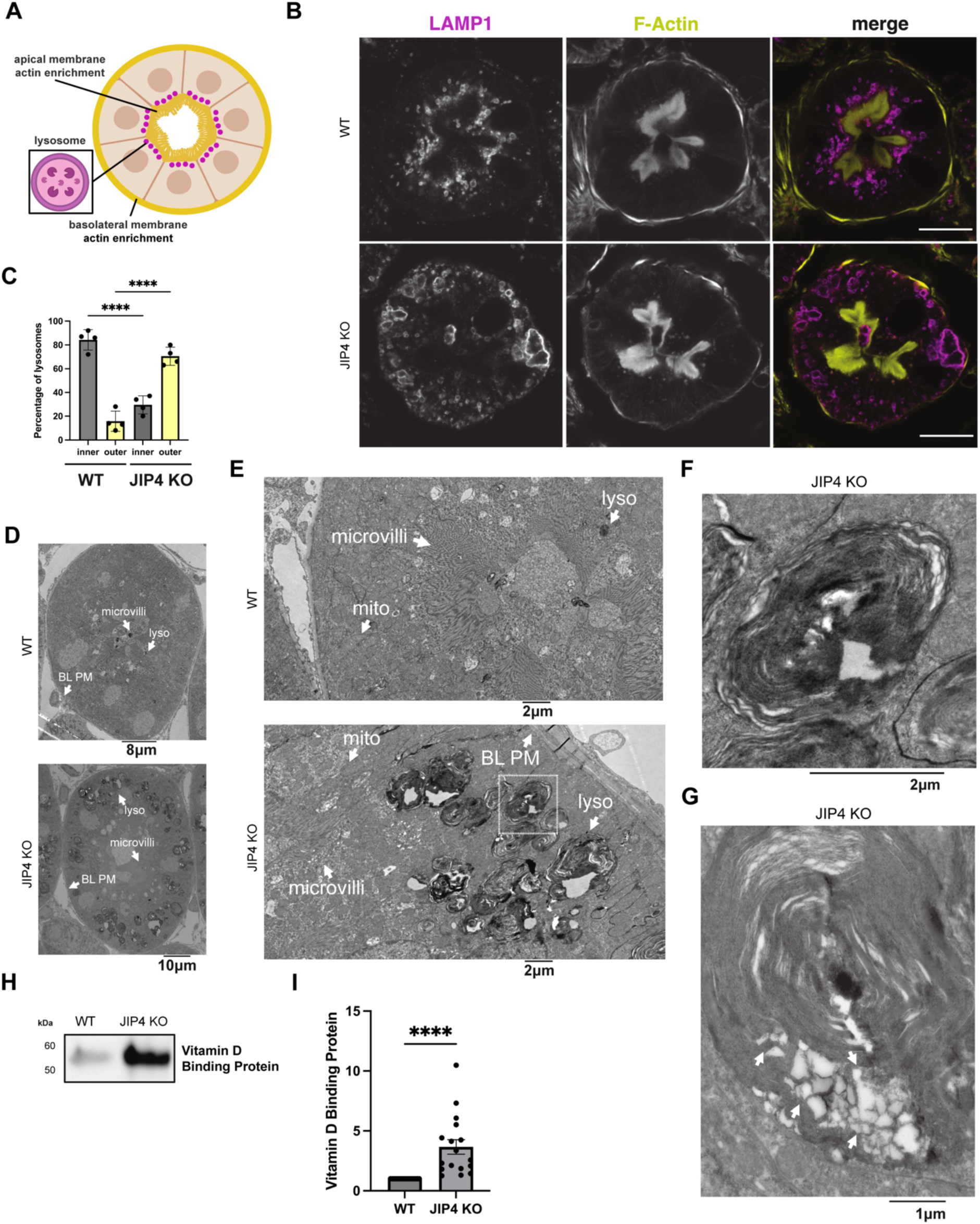
JIP4 KO proximal tubular lysosomes are mislocalized towards the basolateral membrane and contain storage material. **a,** Schematic depicting lysosome enrichment under the apical membrane in a cross-section of a renal proximal tubule. **b,** Immunofluorescent imaging of LAMP1 and F-Actin in WT and JIP4 KO proximal tubules. Scale bar = 20 µm. **c,** Quantification of lysosome localization closer to the apical membrane (inner lysosomes) or basolateral membrane (outer lysosomes) in WT and JIP4 KO proximal tubules in a 14-month-old mouse (mean ± SEM, n=3 biological replicates, ****P<0.0001, one-way ANOVA with Tukey’s multiple comparisons test). **d,** Electron micrograph of WT and JIP4 KO proximal tubule cross sections. Lyso = lysosome, microvilli = apical plasma membrane microvilli, BL PM = basolateral plasma membrane. **e,** Electron micrograph of WT and JIP4 KO proximal tubular lysosomes at either the apical or basolateral membrane, respectively. Mito = mitochondria. **f,** Inset of JIP4 KO electron micrograph in **e. g,** Electron micrograph depicting contents of a JIP4 KO lysosome. **h,** Western blot analysis of vitamin D binding proteins levels in WT and JIP4 KO mouse urine from littermates at 11 months of age. **i,** Quantification of WT and JIP4 KO mouse urine from littermates between 6-14 months of age (mean ± SEM, n=17 biological replicates each, ****P<0.0001, two-tailed Wilcoxan matched-pairs signed rank test)

In addition to their basolateral redistribution, JIP4 KO renal proximal tubule lysosomes are large, and a subset of them forms non-spherical shapes, suggesting a lysosomal storage defect like that found in CTNS KO mouse kidneys (Fig. 5b,c; Cherqui et al., 2002). Confocal microscopy further revealed JIP4 KO autofluorescence within lysosomes in a typical lipofuscin-like pattern, with high levels of autofluorescence when excited by a 568 nm wavelength laser (Extended Data Fig. 8a; Seehafer & Pearce, 2006). Histochemical staining with hemotoxylin and eosin also exhibited eosinophilic abnormalities in JIP4 KO proximal tubules resembling enlarged lysosomes (Extended Data Fig. 8b). However, overall lysosomal protein levels did not change, pointing to storage defects, rather than lysosomal biogenesis, as the cause of lysosomal expansion (Extended Data Fig. 8c). Storage defects in the JIP4 KO mouse kidney were also explored with electron microscopy which revealed that the large, misshapen lysosomes contained lipid storage material along with electron-lucent polygons resembling cystine crystals found in CTNS KO mice (Fig. 5e-g; Cherqui et al., 2002). Analysis of JIP4 KO mouse urine along with WT littermates revealed proteinuria, indicating a failure of proximal tubules to remove protein from urine and mirroring the phenotype of CTNS KO mice (Fig. 5h,I; Cherqui et al., 2002). Further analysis of 2-month-old JIP4 KO mouse proximal tubules revealed that both relocalization of lysosomes to the basolateral membrane and storage defects occur at an early age (Extended Data Fig. 9a-c). Together, these data support an *in vivo* role for JIP4 in maintaining lysosome function that parallels past observations of similar phenotypes in CTNS KO mice.

## Discussion

Our identification of JIP4 as a critical regulator of CTNS stability establishes a novel mechanism for regulation of lysosomal nutrient efflux and, more broadly, cellular cysteine/cystine homeostasis. Our data supports a model wherein JIP4 protects CTNS by limiting the ability of TMEM55B to promote CTNS ubiquitylation and degradation. Major consequences of JIP4 depletion match phenotypes arising from CTNS deficiency.

Although first identified as a scaffold for p38 mitogen-activated protein kinase (MAPK), JIP4 has since been shown to regulate the balance between dynein and kinesin-dependent movement of late endosomes and lysosomes (Kelkar et al., 2005; Montagnac et al., 2009; Willett et al., 2017; Gowrishankar et al., 2021; Celestino et al., 2022; Cason & Holzbaur, 2023). Reduced coupling of lysosomes to active dynein-dynactin explains peripheral accumulations of lysosomes in JIP4 KO cells (Fig. 5b,c; Extended Data Fig. 1; Extended Data Fig. 7b; Extended Data Fig. 9a). The mechanistic basis for such dynein-dynactin activation by JIP4 can be inferred from recently defined structure and biochemistry of dynein-dynactin activation by the closely related, neuronally enriched, JIP3 protein (Singh et al., 2024). The JIP4-TMEM55B interaction was previously proposed to recruit JIP4 to lysosomes to promote this dynein-dynactin-dependent movement (Willett et al., 2017). Importantly, our data supports a separate function for JIP4 and TMEM55B in the regulation of CTNS-dependent efflux of cystine from lysosomes. Our results thus establish coordination between lysosomal metabolism and lysosome movement through dual action of JIP4 in connecting lysosomes to molecular motors as well as suppressing turnover of CTNS.

The dual roles of JIP4 are evident in the renal proximal tubule, where the endolysosomal system is adapted to support retrieval of proteins and other nutrients from the glomerular filtrate (Rodman et al., 1986; Eshbach & Weisz, 2017). To this end, lysosomes are highly concentrated under the apical plasma membrane of the proximal tubule, and JIP4 localizes to these lysosomes (Fig. 5a-c; Extended Data Fig. 7a,b; Extended Data Fig. 9a). However, in JIP4 KO mouse renal proximal tubules, lysosomes are redistributed towards the basolateral membrane, consistent with a defect in dynein-dynactin directed movement. In addition to being mislocalized, these JIP4 KO proximal tubular lysosomes are strikingly enlarged and develop signs of severe lysosomal storage (Fig. 5d-f; Extended Data Fig. 8; Extended Data Fig. 9b). These results illustrate an *in vivo* relationship between JIP4-dependent lysosome positioning and degradative functions.

Our new observations for JIP4-mediated control of CTNS have parallels to the control of budding yeast Ypq1 protein. Ypq1, a member of the PQ-loop transporter family that also contains CTNS, functions at the yeast vacuole and undergoes ubiquitylation and ESCRT-mediated degradation in response to starvation of its substrate lysine (Li et al., 2015; Zhu et al., 2017; Arines et al., 2021). This system, which incorporates the transmembrane adaptor Ssh4 that directs Ypq1 ubiquitylation by the NEDD4 family E3 ligase Rsp5, has parallels to the JIP4-TMEM55B-CTNS system. Post-translational control of lysosomal transporters may allow cells to rapidly adapt to changes in cellular demand for nutrients (Prus et al., 2024). By defining JIP4 as a critical upstream regulator of CTNS, we provide new insight into how mammalian cells modulate lysosomal transporter levels to match metabolic demand.

Our work establishes that CTNS is subject to a regulated degradation that controls its activity on lysosomes. Controlled cystine release acts as a bulwark against oxidative stress while also preventing mitochondrial toxicity due to cysteine accumulation in the cytosol, as precise cysteine compartmentalization within lysosomes is critical for proper mitochondrial function, prevention of ferroptosis and management of oxidative stress (Hughes et al., 2020; Armenta et al., 2022; He et al., 2023; Swanda et al., 2023). While cystinosis is classically attributed to inactivating mutations in CTNS, our data raise the possibility that defects in CTNS regulation, such as impaired JIP4 function, could similarly disrupt cystine export and contribute to mismanagement of cysteine in disease.

Our new findings have clinical relevance with respect to recently reported JIP4 loss-of-function mutations in humans. Homozygous recessive mutations in JIP4 were very recently shown to result in neurodevelopmental impairment, motor delay, hypopigmentation, and cataracts (Acosta-Baena et al., 2024; Alfadhel et al., 2025). The pigmentation and ocular symptoms arising from JIP4 mutations are reminiscent of cystinosis (Gahl et al., 2002; Elmonem et al., 2016). We thus hypothesize that human JIP4 loss-of-function mutations may impair CTNS regulation and predispose patients to cystinosis-like phenotypes, even in the presence of an intact CTNS gene. This raises the possibility that cysteamine treatment may benefit patients with JIP4 mutations. Additionally, JIP4 and TMEM55B have been reported to function downstream of leucine rich repeat kinase 2 (LRRK2) at lysosomes (Bonet-Ponce et al., 2020; Pal et al., 2023). Aberrant JIP4-TMEM55B-dependent control of lysosomal cystine efflux may thus contribute to the effects of LRRK2 gain-of-function mutations that give rise to Parkinson’s disease (Alessi & Pfeffer, 2024). Collectively, this study establishes a novel pathway for JIP4-dependent control of lysosomal cystine homeostasis with a broad range of potential physiological and pathophysiological impacts.

## Materials and Methods

### Cell culture and transfection

HeLa-M cells (kindly provided by Pietro De Camilli, Yale University) were maintained in DMEM + 10% FBS + 1% penicillin/streptomycin (Thermo Fisher Scientific) at 37°C with 5% CO_2_. Experiments to test vacuole formation used RPMI media without amino acids (US Biological) supplemented with minimal essential amino acids (Gibco) +L-glutamine (4mM, Gibco). The MEM–amino acids solution at 1× contains L-arginine (600 µM), L-cystine (100 µM), L-histidine (200 µM), L-isoleucine (400 µM), L-leucine (400 µM), L-lysine (396 µM), L-methionine (101 µM), L-phenylalanine (200 µM), L-threonine (400 µM), L-tryptophan (50 µM), L-tyrosine (199 µM), and L-valine (400 µM). Dialyzed serum was used where indicated (Gibco). For cysteamine experiments, cysteamine (Sigma-Aldrich) was added to starvation media at 1 mM final concentration for 1 h before lysis.

For plasmid DNA transfections, 1-3 x 10^5^ cells were plated per well in a 6-well dish. The next day, transfections were performed with 2ug of plasmid DNA, 2uL of Lipofectamine 2000 (Invitrogen), and 100uL Opti-MEM (Invitrogen). For transfections of smaller or larger dishes, volumes were scaled proportionally. A detailed cell culture protocol can be accessed at https://doi.org/10.17504/protocols.io.8epv5rrq4g1b/v1

### CRISPR/Cas9 genome editing

gRNA targeting exon 1 of *CTNS* were cloned into the px459 plasmid, as previously described (Amick et al., 2020). 2µg of plasmid DNA was transfected with Lipofectamine 2000 into 250,000 HeLa cells in a 6-well dish. The next day, transfected cells were selected with 1.25 µg/ml puromycin for 3 days. Surviving cells were subsequently plated at clonal density. Following the selection and expansion of colonies, CTNS KOs were identified by sequencing of PCR-amplified genomic DNA. To sequence genomic DNA, it was extracted (QuickExtract DNA extraction solution; Epicentre Biotechnologies), and the region of interest was amplified by PCR (primers described in Table S1), cloned into the pCR-Blunt TOPO vector (Zero Blunt TOPO PCR cloning kit; Thermo Fisher Scientific), and transformed into TOP10-competent *E. coli* cells. Plasmid DNA was then isolated from colonies and sequenced to define the genotype of the locus of interest. CTNS KO clones were additionally verified through Synthego ICE analysis of sequencing results (https://ice.editco.bio/#/). The method used for CRISPR/Cas9 genome editing to insert the 2xHA epitope tag at the endogenous CTNS locus was described previously (Amick et al., 2018). The single-strand DNA oligonucleotide homology-directed repair donor template was designed with asymmetric homology arms on the protospacer-adjacent motif proximal and distal sides of the cut site to enhance the efficiency of tag insertion. Clonal cell populations were isolated and screened for HA signal by immunoblotting and immunofluorescence. To further confirm the correct in-frame insertion of the 2xHA tag in these cells, genomic DNA surrounding the site of tag insertion was PCR amplified and sequenced as described above. The method used for CRISPR/Cas9 genome editing to insert the GFP epitope tag at the endogenous LAMP locus was described previously (Boecker et al., 2020). ssDNA and crRNAs were ordered through IDT.

### Stable cell line generation

A lentiviral strategy was used for stable cell line generation of CTNS-FLAG. Briefly, 2.5 x 10^5^ HEK293T cells were plated per well in a 6-well dish coated with poly-D-lysine (Sigma-Aldrich). The following day, cells were transfected using Lipofectamine 2000 (Invitrogen), psPAX2, pCMV-VSVG, and pLVX-Puro-CTNS-FLAG plasmids. psPAX2 was a gift from Didier Trono, École polytechnique fédérale de Lausanne, Lausanne, Switzerland (Addgene plasmid #12260); pCMV-VSV-G was a gift from Bob Weinberg, Whitehead Institute for Biomedical Research, Cambridge, MA (Addgene plasmid #8454). After 48 h, the viral supernatant was filtered through a 0.45-μm filter (Pall Corporation) onto HeLa cells to be transduced. Polybrene solution (8 μg/ml; EMD Millipore) was added to increase uptake of virus. 2 μg/ml puromycin (Thermo Fisher Scientific, Gibco) was then used to select for stable transductant cells. Clonal cell populations were isolated and screened for FLAG signal by immunoblotting and immunofluorescence. JIP4 KO CTNS-FLAG and JIP4 KO CTNS-FLAG-K->A were created by adding JIP4 gRNAs to each clonal line. Clonal lines were confirmed via immunoblotting and immunofluorescence.

### Plasmids

CTNS-FLAG plasmids were generated as follows. The CTNS coding sequence followed by a FLAG tag (DYKDDDDK tag) and a stop codon was purchased as synthetic double stranded DNA (gBlock, Integrated DNA Technologies). For lentivirus-mediated transgenic expression, this DNA was inserted into SmaI-digested pLVX vector (Clontech) by Gibson Assembly (NEBuilder HiFi DNA Assembly; New England BioLabs) according to manufacturer protocols. The synthetic CTNS-RUSH coding sequence was also purchased as a gBlock from Integrated DNA Technologies. This DNA was inserted into SmaI-digested pEGFPN2 vector (Clontech) by Gibson Assembly (NEBuilder HiFi DNA Assembly; New England BioLabs) according to manufacturer protocols. CTNS K->A mutations were generated by site-directed mutagenesis in combination with fragment assembly using Gibson Assembly (NEBuilder HiFi DNA Assembly; New England BioLabs) and the “Improved Methods for Site-directed Mutagenesis using NEBuilder® HiFi DNA Assembly Master Mix” protocol from NEB. Stbl3 *E. coli* (Invitrogen) was used for cloning. PiggyBac-SKIP-myc was purchased from VectorBuilder. mCherry-TMEM55B plasmid was kindly provided by Pietro De Camilli. All plasmids were sequence verified.

### Immunoprecipitation and immunoblotting

For cell lysis, one 85% confluent 15-cm plate was used per sample. Cells were washed 2X with PBS, scraped in ice-cold lysis buffer consisting of 50 mM Tris, pH 7.4, 150 mM NaCl, 1 mM EDTA, and 1% Triton X-100 plus protease and phosphatase inhibitor cocktails (Complete Mini, EDTA-free, PhosSTOP; Roche). Lysates were incubated on ice for 5 minutes and then centrifuged at 14,000 RPM (4°C) for 8 minutes. Protein concentrations were measured using Coomassie Plus Protein Assay Reagent (ThermoFisher Scientific, 23236) as per manufacturer’s protocol. Whole cell lysates were processed in a second tube, and an equivalent volume of 4x Laemmli Buffer was added to each lysate. These samples were supplemented with 6.187% fresh B-mercaptoethanol (Sigma-Aldrich) and heated at 42°C for 3 minutes. Immunopreciptation lysates were then immunoprecipitated using 12uL anti-FLAG M2 affinity gel (Sigma-Aldrich) or anti-RFP beads (Rockland) that were pre-washed 3X with 0.1% Triton lysis buffer. The same amount of protein was used for each sample. Where needed, samples were supplemented with lysis buffer to maintain the same protein concentration and volume. Lysates were incubated with beads rotating end-over-end for 1 hour at 4°C. Resin was subsequently washed five times with lysis buffer and then eluted with 2x Laemmli buffer at 42°C for 3 minutes. The sample was then transferred to another microcentrifuge tube. Immunoblotting was performed with 4–15% gradient Mini-PROTEAN TGX precast polyacrylamide gels and nitrocellulose membranes (Bio-Rad). Blots were blocked with 5% nonfat dry milk (AmericanBIO), and antibodies were incubated with 5% nonfat dry milk or BSA (AmericanBIO) in TBS with 0.1% Tween. Chemiluminescence detection of HRP signals from secondary antibodies was performed on a Chemi-Doc imaging station (Bio-Rad). A more detailed protocol can be accessed at dx.doi.org/10.17504/protocols.io.5qpvo9bmdv4o/v1

### siRNA

siRNA-mediated knockdowns of target gene expression were accomplished using Dharmacon siGENOME pooled siRNAs. siRNA transfections were performed with 2.5uL of 20uM TMEM55B along with 2.5uL TMEM55A siRNA (On-Target Plus Smartpool, Dharmacon) or non-targeting siRNA (Dharmacon), 5uL Lipofectamine RNAiMAX transfection reagent (Invitrogen), and 200uL OptiMEM (Invitrogen) as per the manufacturer’s protocols using 100 nM siRNA pool. After 48 hours, cells were lysed and subjected to immunoblotting. A more detailed protocol can be accessed at: dx.doi.org/10.17504/protocols.io.4r3l29owjv1y/v1

### Immunofluorescence and imaging

Cells were fixed in a 4% paraformaldehyde (Electron Microscopy Sciences, 19202)/sodium phosphate buffer (pH 7.3 / Buffer: 153.56 mM Sodium phosphate, dibasic, anhydrous, J.T. Baker 3828, 53.63 mM Sodium dihydrogen phosphate monohydrate, J.T. Baker 3818) for 30 minutes at room temperature. Cells were washed 3 times for 5 minutes each with PBS. Cells were permeabilized by immersing coverslips in ice cold methanol for 3 seconds and followed by PBS rinses. Cells were blocked in 3% BSA in PBS. Primary antibody was added overnight at 4°C. Cells were washed 3 times for 5 minutes each with PBS. Secondary antibody was added for 1 hour at room temperature in the dark. Cells were washed 3 times for 5 minutes each with PBS. Coverslips were mounted onto microscope slides (Thermo Fisher Scientific, 12-550-143) with Prolong Gold mounting media (Thermo Fisher Scientific, P36935) and stored at 4°C. Imaging was performed on a Zeiss LSM880 confocal laser scanning microscope with Airyscan using a Plan Apochromat 63× objective (NA 1.4), a 32-channel gallium arsenide phosphide photomultiplier tube detector, and 488-, 561-, and 633-nm laser lines. Images were processed in FIJI. Quantifications of colocalization between CTNS-FLAG and LAMP1 were performed using the Coloc2 plugin in FIJI, plotting Mander’s coefficient after adjusting for threshold. Lysosome positioning quantifications in cells and in kidneys were performed using separate, but similar, CellProfiler analyses measuring the numbers of lysosomes within 10 concentric rings. Rings 1-5 were named “inner lysosomes,” while rings 6-10 were named “outer lysosomes”

### RUSH assay

Cells were plated at 2 x 10^5^ on clean coverslips in a 24 well plate. After one day, cells were transfected with 20µL OptiMEM, 0.4ug CTNS-RUSH plasmid, and 0.4µL Lipofectamine 2000. After one additional day, 40 µM biotin (EMD Millipore) diluted in water and 178 µM cycloheximide (Invitrogen) diluted in DMSO were added simultaneously to cells for 1, 2, or 4 hours. After the indicated amount of time, cells were fixed and processed as described above for immunofluorescence and imaging.

### Live imaging

HeLa cells were plated at 100,000 cells in 2 ml medium onto 35-mm MatTek glass-bottom dishes 1 day before imaging. Live-cell imaging was performed in an environment-controlled chamber set at 37°C and 5% CO_2_ on a Zeiss LSM880 confocal laser scanning microscope with Airyscan as previously described.

### Metabolite profiling

To study metabolic changes associated with vacuole formation, cells were seeded at 5 x 10^5^ in 15 cm^2^ plates for 72 hr prior to harvesting. To prepare cellular metabolite extracts, cells were washed twice in 1ml ice cold PBS and then scraped in 500 µl acetonitrile-methanol-water (27:9:1 vol/vol/vol). The extraction mixture was vortexed and centrifuged at 19,000 × *g* for 20 min at 4 °C, and 150 μl supernatants were transferred into LC-MS glass vials for analysis.

LC/MS-based analyses were performed on a Q Exactive Plus benchtop Orbitrap mass spectrometer equipped with an Ion Max source and a HESI II probe, which was coupled to a Vanquish UHPLC. Polar metabolite detection method was adapted with minor modification from previous literature (Shen et al., 2017; Shi et al., 2022).

Polar metabolites were analyzed on Xbridge BEH Amide XP HILIC Column, 100 Å, 2.5 μm, 2.1 mmx100 mm (Waters, 186006091) for chromatographic separation. The column oven temperature was 27 °C and the autosampler was 4 °C. Mobile phase A: 5% acetonitrile, 20 mM ammonium acetate/ammonium hydroxide, pH 9, and mobile phase B: 100% acetonitrile. LC gradient conditions at a flow rate of 0.220 ml/min as follows: 0 min: 85% B, 0.5 min: 85% B, 9 min: 35% B, 11 min: 2% B, 13.5 min: 85% B, 20 min: 85% B. The mass data were acquired in the polarity switching mode with full scan mode in a range of 70–1000 m/z, with the resolution at 70,000, the AGC target at 3e^6^, and the maximum injection time at 80 ms, the sheath gas flow at 50 units, the auxiliary gas flow at 10 units, the sweep gas flow at 2 units, the spray voltage at 2.5 kV(-) and 3.8 kV(+), the capillary temperature at 310 °C, and the auxiliary gas heater temperature at 370 °C. Compound discoverer (Thermo Fisher Scientific) was used for peak picking and intensity.

### Mutant Mouse Generation

JIP4 KO mutant mice were generated by the Yale Genome Editing Center. C57Bl/6J zygotes were electroporated with a pair of Cas9/sgRNA ribonucleoprotein complexes (sgRNA1: TAGTAAGGGCTACTGTAGTG; sgRNA2: TGGATGTGCAAATAACGGAG) that targeted sequences flanking exon 4. This resulted in a deletion of 3180 base pairs that included the 95 base pairs encoded by exon 2 (Extended Data Fig. 6).

### Immunohistochemistry of Mouse Kidneys

12–14-month-old (Fig. 5, Extended Data Fig. 7-8) or 2-month-old mice (Extended Data Fig. 9) were anesthetized with isoflurane and transcardially perfused with ice-cold PBS followed by 4% PFA in PBS. Kidneys were dissected and post-fixed in 4% PFA at 4°C overnight. Kidneys were then rinsed in PBS and embedded in 20% sucrose solution overnight and then kept in 30% sucrose for storage. Kidneys were frozen in OCT (Fisher Healthcare 23-730-571) at −80°C, and sections (40-50 µm) were cut using a cryostat. Fixed sections were permeabilized with 0.3% Triton X-100 in PBS for 1 hour at room temperature and blocked with 5% normal donkey serum (NDS) in PBS for 2 hours at room temperature. Sections were incubated overnight at 4°C with primary antibodies diluted in PBS containing 1% NDS and 0.1% Triton X-100. After washing with PBS, sections were incubated with appropriate fluorophore-conjugated secondary antibodies for 2 hours at room temperature in the dark. Sections were mounted on glass slides using Fluoromount-G (SouthernBiotech), and images were captured using a Zeiss LSM880 confocal laser scanning microscope with Airyscan as described for immunofluorescence analysis.

### Proteinuria analysis

Urine from WT, heterozygous, and JIP4 KO littermates was mixed with an equal volume of Laemmli Buffer + 6.125% beta-mercaptoethanol and heated for 3 minutes at 95°C and processed for immunoblotting.

### Transmission Electron Microscopy

Isolated kidneys were fixed in 2% paraformaldehyde in 0.1 M sodium cacodylate buffer pH 7.4 containing 2% sucrose for 1 h and post fixed in 1% osmium tetroxide for 1 h. The sample was rinsed in buffer and en-bloc stained in aqueous 2% uranyl acetate for 1 h followed by rinsing in distilled water, dehydrated in an ethanol series, and infiltrated with Embed 812 (Electron Microscopy Sciences) resin. The samples were placed in silicone molds and baked at 60 °C for 24 h. Hardened blocked were sectioned using a Leica UltraCut UC7. Sixty-nanometer sections were collected on formvar-coated nickel grids and 250-nm sections on copper slot grids and stained using 2% uranyl acetate and lead citrate. Samples were viewed on an FEI Tencai Biotwin transmission electron microscope at 80 kV. Images were taken using a Morada CCD and iTEM (Olympus) cellSens Dimension software.

### Statistical analysis

Statistical analysis was performed with Prism 10 software, with specific details about the statistical tests conducted, the number of independent experiments, and P values provided in the corresponding figure legends.

## Acknowledgments

We are grateful to all members of the Ferguson laboratory, Pietro De Camilli, and Michael Caplan for helpful advice and discussions. We thank Joseph Amick for critical readings of early manuscript drafts. This research was supported by grants from the National Institutes of Health (AG085824 and AG062210) as well as from Aligning Science Across Parkinson’s disease (ASAP-000580) through the Michael J. Fox Foundation for Parkinson’s Research (MJFF) to S.F.; and NIGMS (R35GM150619) to H.S. We are grateful to Suxia Bai and Timothy Nottoli at the Yale Genome Editing Center for generation of the JIP4 mutant mice and to Taylor Skibitcky for help with mouse genotyping. We thank Berrak Ugur (Yale) and Benjamin Johnson (Yale) for managing our ASAP project. We thank the Yale Center for Cellular and Molecular Imaging’s Electron Microscopy Core facility for electron microscopy of mouse kidneys. We thank Biorender for use in illustrations. LMN was supported by the National Science Foundation Graduate Research Fellowship. The authors declare no competing financial interests.

## Author Contributions

L.M.N. and S.F. conceptualized the project. L.M.N. performed most of the experiments. X.S. and H.F. performed metabolite profiling via mass spectrometry. A.F. created JIP4 KO and rescue cell lines, validated reagents, and made key observations that were critical for initiation of the project. All authors edited and provided comments on the manuscript.

## Availability Statement

The data, protocols, and key lab materials used and generated in this study are listed in a Key Resource Table alongside their persistent identifiers at 10.5281/zenodo.15588218. No new code was generated for this study; all data cleaning, preprocessing, analysis, and visualization was performed using FIJI/ImageJ, Alphafold, ChimeraX, CellProfiler, and GraphPad Prism.

---

**Extended Data Fig. 1:**
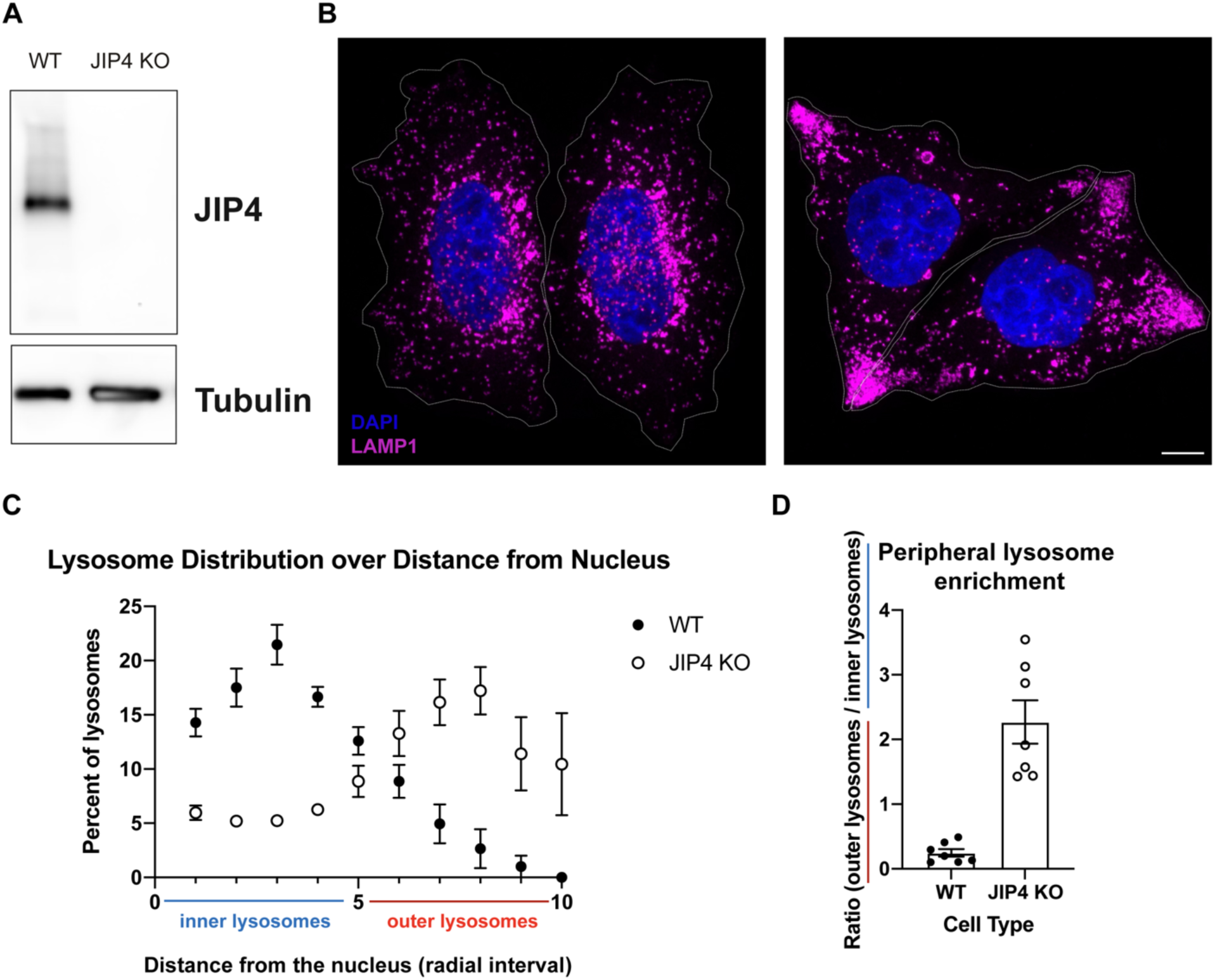
JIP4 KO lysosomes cluster at the periphery of HeLa cells, related to Fig. 1. **a,** Western blot analysis of WT and JIP4 KO cells. **b,** Immunofluorescent imaging of WT and JIP4 KO cells. Scale bar = 10 µm. **c,** Quantification of WT and JIP4 KO lysosomes by radial interval. **d,** ratio of inner vs outer lysosomes for WT vs JIP4 KO cells (n=3 independent experiments, 2-3 cells per experiment).

**Extended Data Fig. 2:**
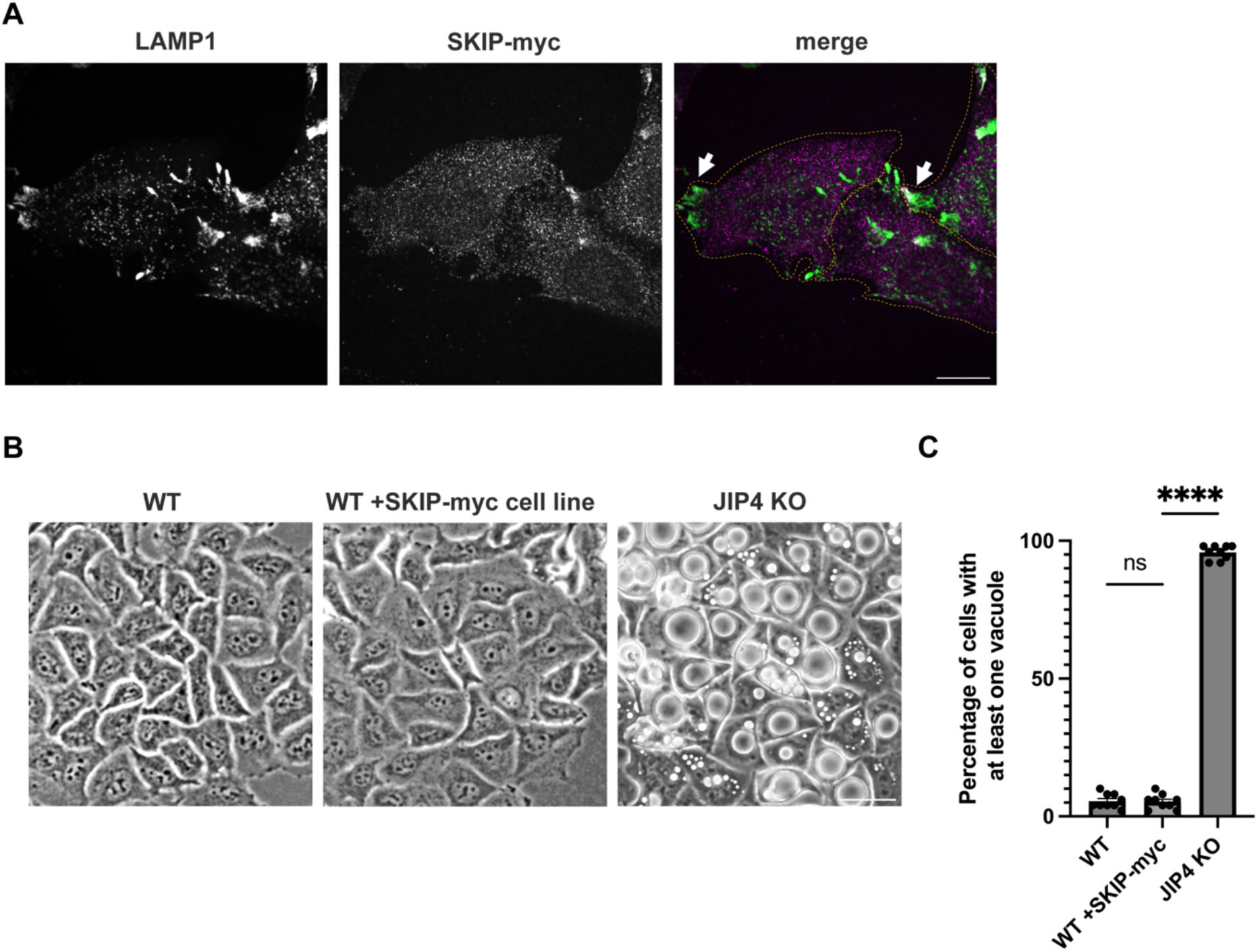
Moving lysosomes to the cellular periphery is not sufficient to create vacuoles, related to Fig. 1. **a,** Immunofluorescent imaging of WT HeLa cells with stably expressed +SKIP-myc. Scale bar = 20 µm **b,** Brightfield imaging of WT, WT +SKIP-myc, and JIP4 KO 3 days post-plating. Scale bar = 50 µm. **c,** Quantification of WT vs WT +SKIP-myc vs JIP4 KO cells in **b**, (mean ± SEM, n=3 independent experiments, ****P<0.0001, unpaired *t* test).

**Extended Data Fig. 3:**
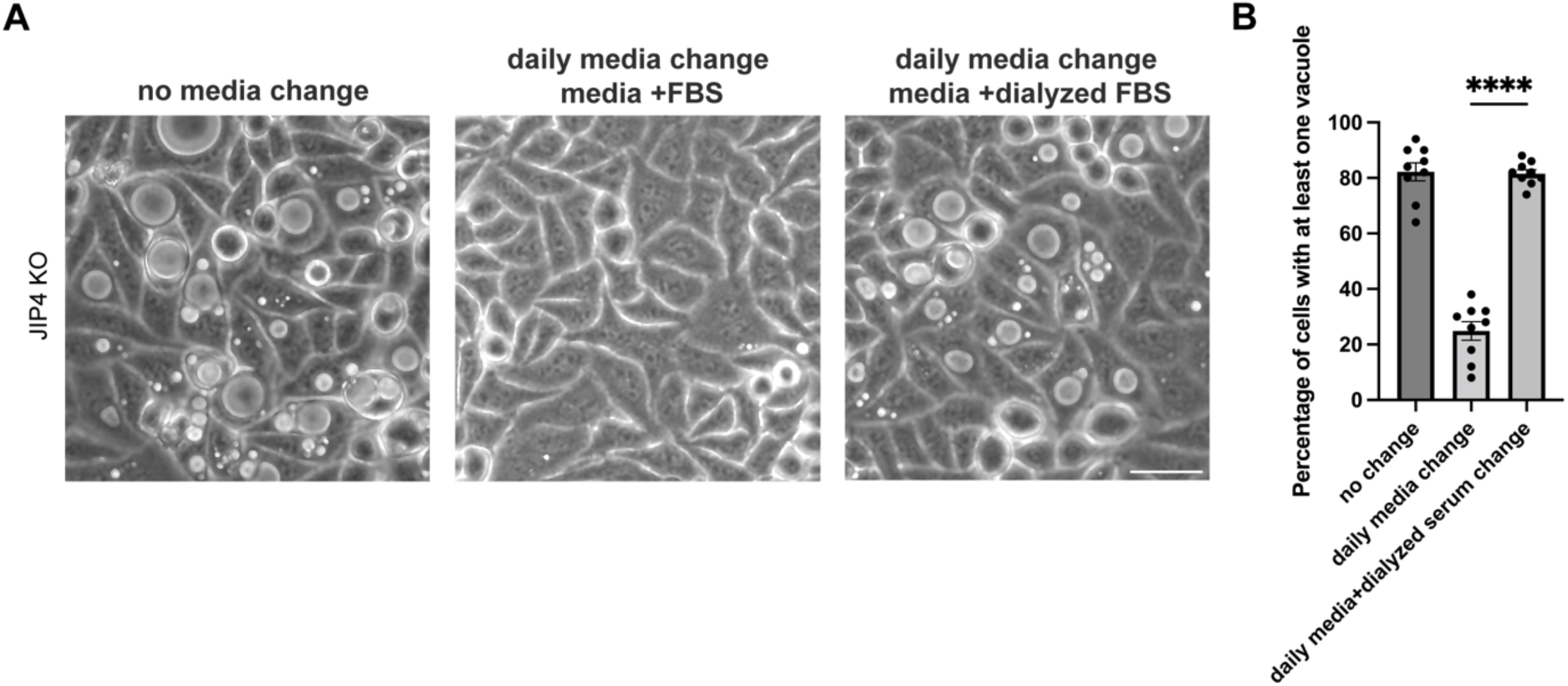
A low molecular weight component of media prevents vacuole formation in JIP4 KO cells, related to Fig. 1. **a,** Brightfield imaging of JIP4 KO cells after three days with no media change, daily media changes, and daily media changes with dialyzed serum. Scale bar = 50 µm.**b,** Quantification of JIP4 KO cells in **a** with at least one vacuole visible by brightfield (mean ± SEM, n=3 independent experiments, ****P<0.0001, one-way ANOVA with Tukey’s multiple comparisons test).

**Extended Data Fig. 4:**
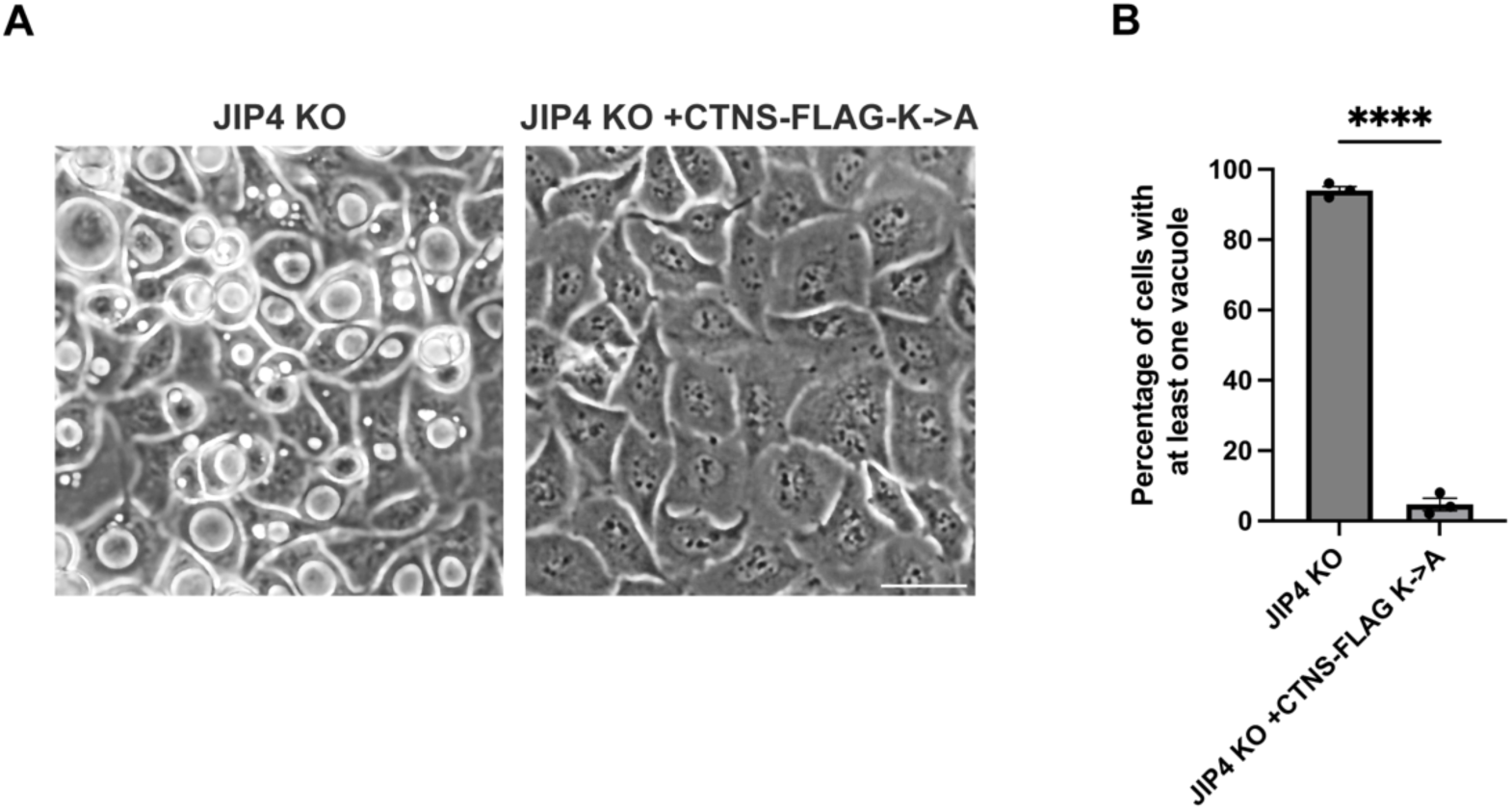
Overexpression of CTNS with K->A mutations prevents vacuoles in JIP4 KO cells, related to Fig. 3. **a,** Brightfield imaging of JIP4 KO and JIP4 KO +CTNS-FLAG-K->A after 3 days post-plating. Scale bar = 50 µm. **b,** Quantification of JIP4 KO vs JIP4 KO +CTNS-FLAG-K->A cells in **a** (mean ± SEM, n=3 independent experiments, ****P<0.0001, unpaired *t* test).

**Extended Data Fig. 5:**
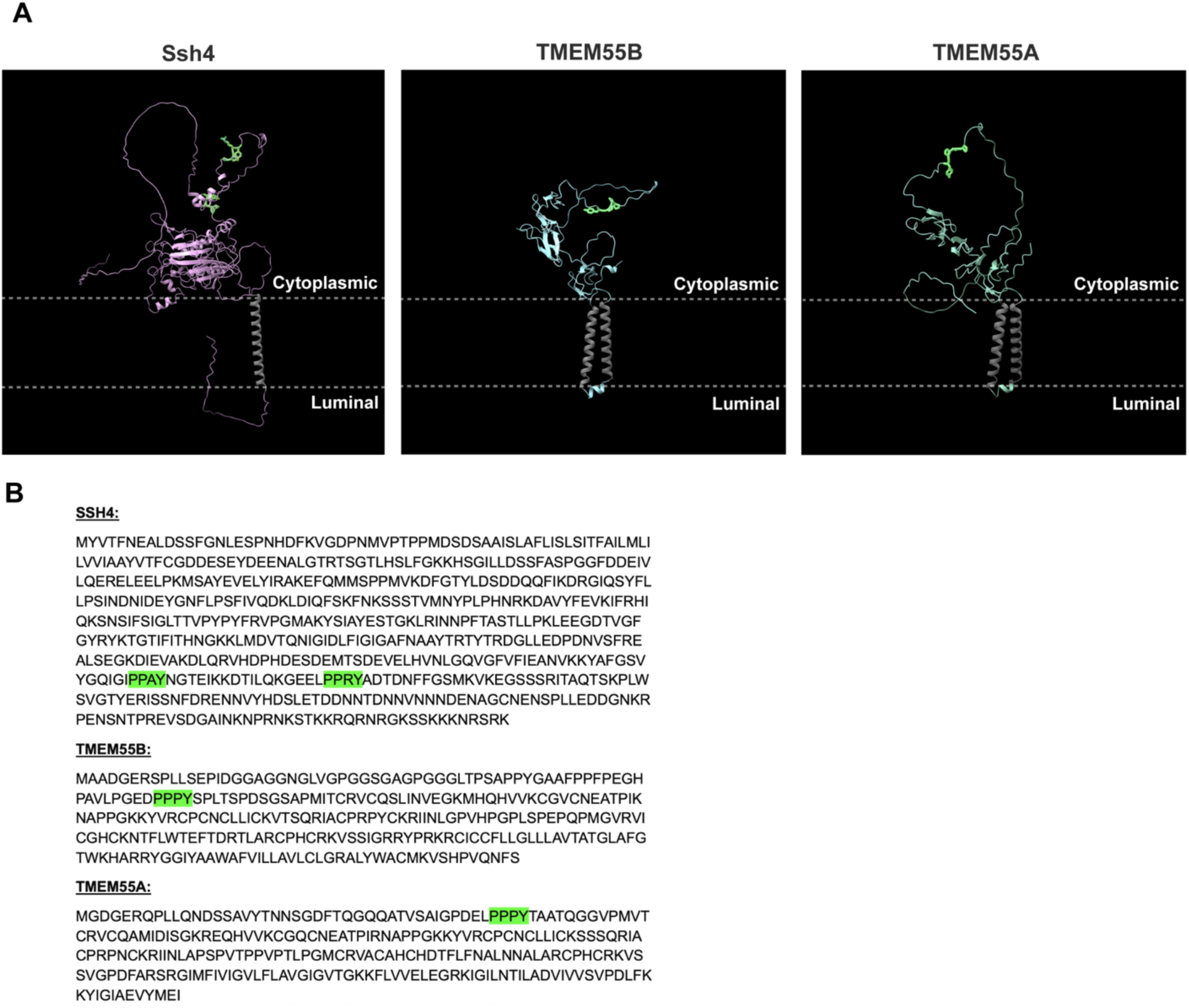
TMEM55A/B and Ssh4 are lysosomal and yeast vacuolar proteins with cytoplasmic-facing PY motifs, related to Fig. 4. **a,** Alphafold models of *Saccharomyces cerevisiaie* Ssh4 and *Homo sapiens* TMEM55B, and TMEM55A proteins. Transmembrane domains colored in gray and PY domains highlighted in green. Cytoplasmic-facing and lysosome luminal-facing regions marked. **b,** Sequences of Ssh4, TMEM55B, and TMEM55A with PY domains highlighted in green.

**Extended Data Fig. 6:**
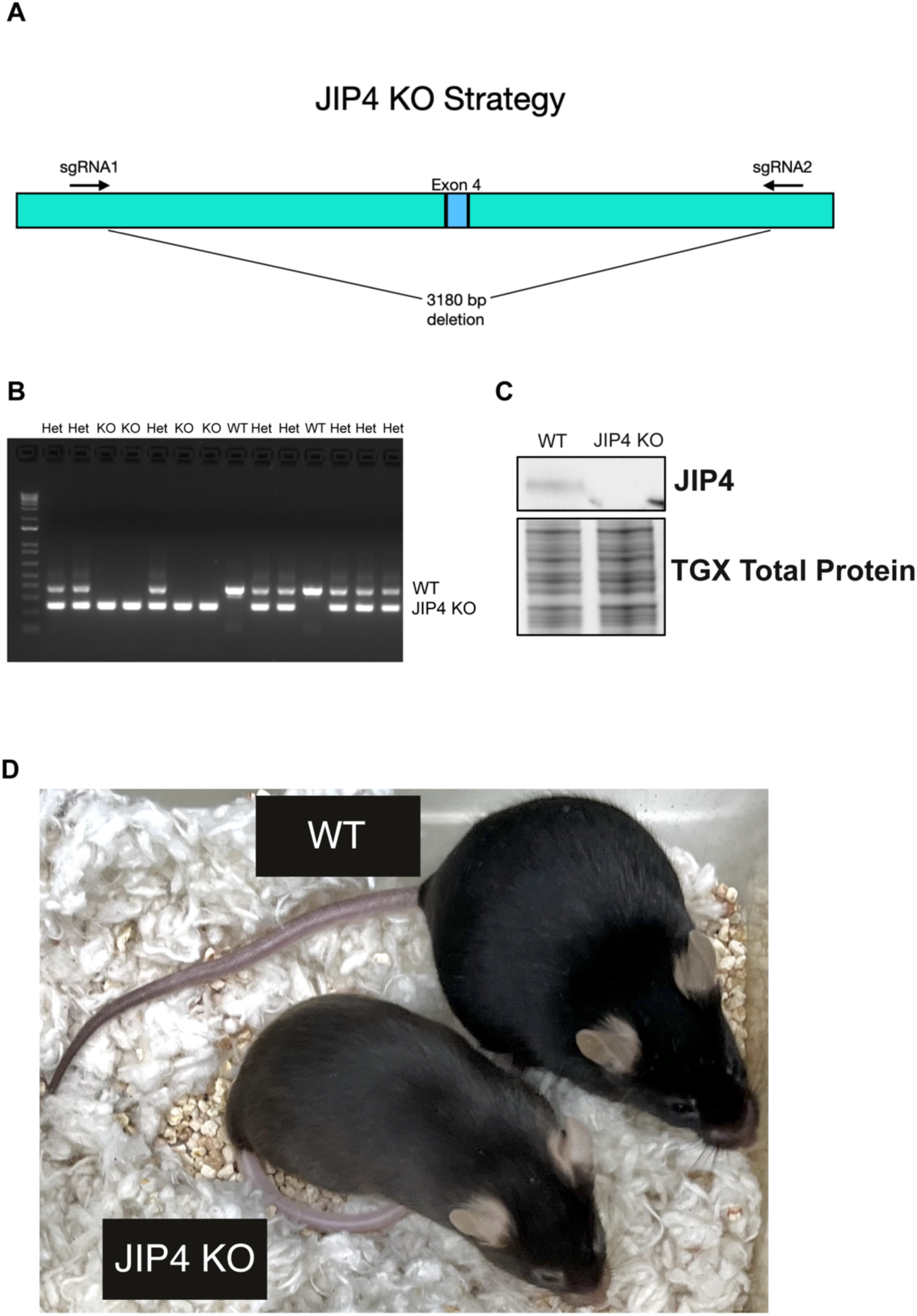
JIP4 KO mice exhibit hypopigmentation, related to Fig. 5. **a,** Schematic of knockout strategy through the deletion of exon 4. **b,** PCR results of 14 mice genotyped from 2 litters. **c,** Western blot analysis of WT and JIP4 KO kidneys. **d,** Images of WT and JIP4 KO littermates.

**Extended Data Fig. 7:**
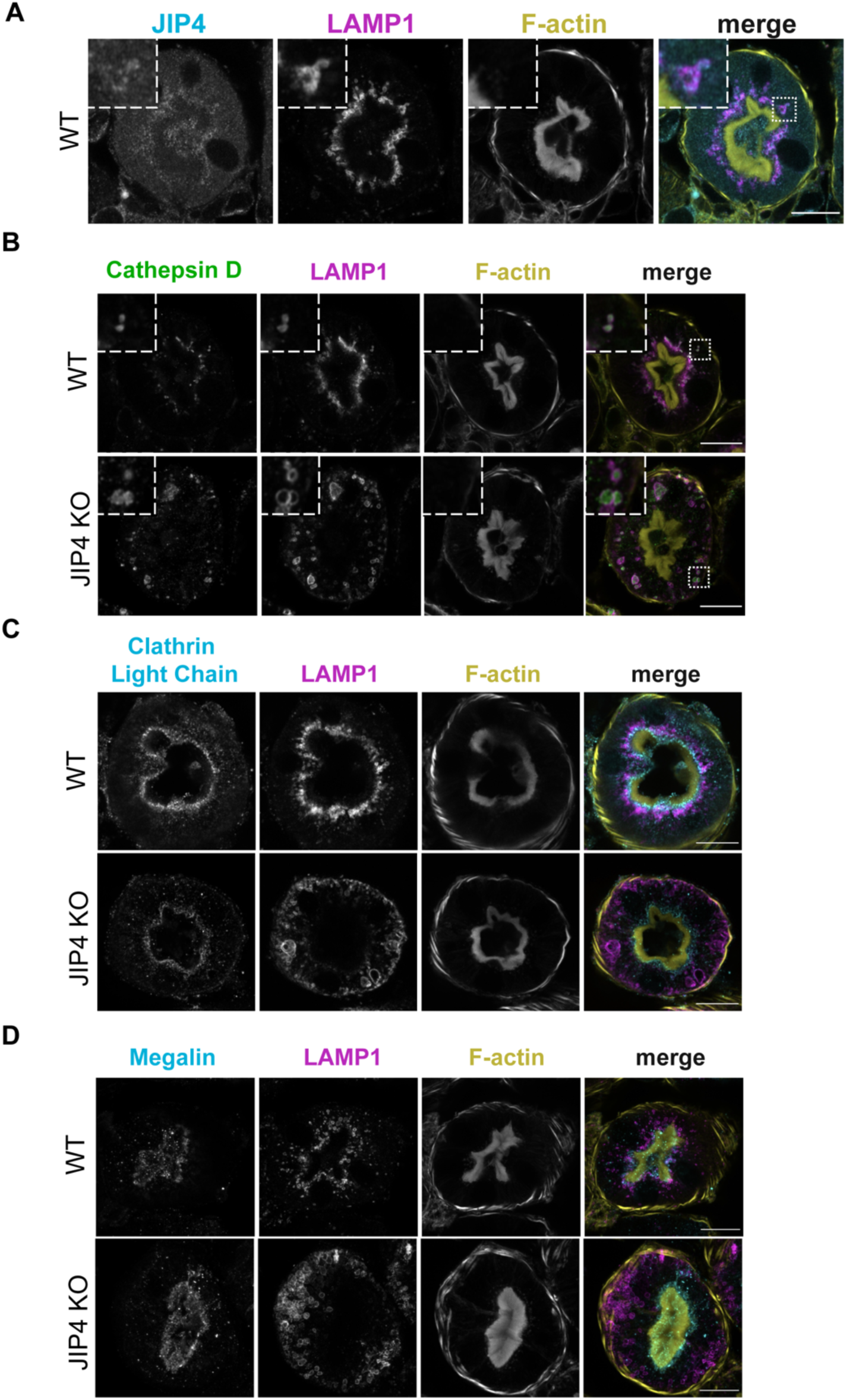
JIP4 is required for apical enrichment of lysosomes in proximal tubules, related to Fig. 5. **a,** JIP4, LAMP1, and F-actin staining in WT mouse renal proximal tubule cross-sections from 12–14-month-old mice. Scale bar = 20 µm. **b,** Cathepsin D, LAMP1, and F-actin staining in WT and JIP4 KO renal proximal tubule cross-sections. Scale bar = 20 µm. **c,** Clathrin Light Chain (CLC), LAMP1, and F-actin staining in WT and JIP4 KO renal proximal tubule cross-sections. Scale bar = 20 µm. **d,** megalin, LAMP1, and F-actin staining in WT and JIP4 KO renal proximal tubule cross-sections. Scale bar = 20 µm.

**Extended Data Fig. 8:**
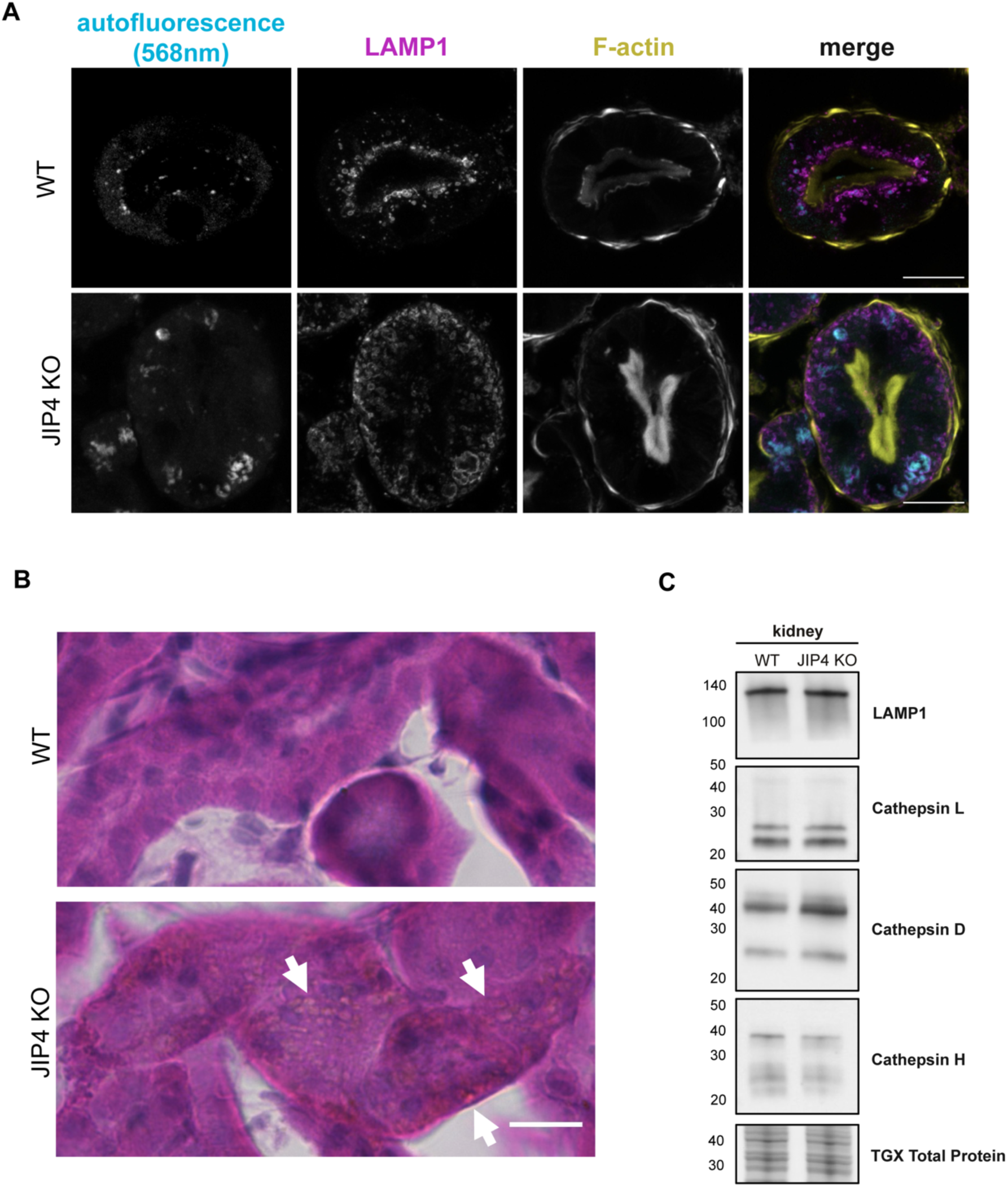
JIP4 KO kidney lysosomes contain storage materials, but lysosome protein levels remain stable, related to Fig. 5. **a,** Lysosome-localized autofluorescence in WT and JIP4 KO kidneys appears when excited with 568nm laser in 12–14-month-old mice. Scale bar = 20 µm. **b,** Hemotoxylin and eosin staining in WT and JIP4 KO proximal tubules. Scale bar = 20 µm. **c,** Western blot analysis of lysosomal proteins in JIP4 KO kidneys.

**Extended Data Fig. 9:**
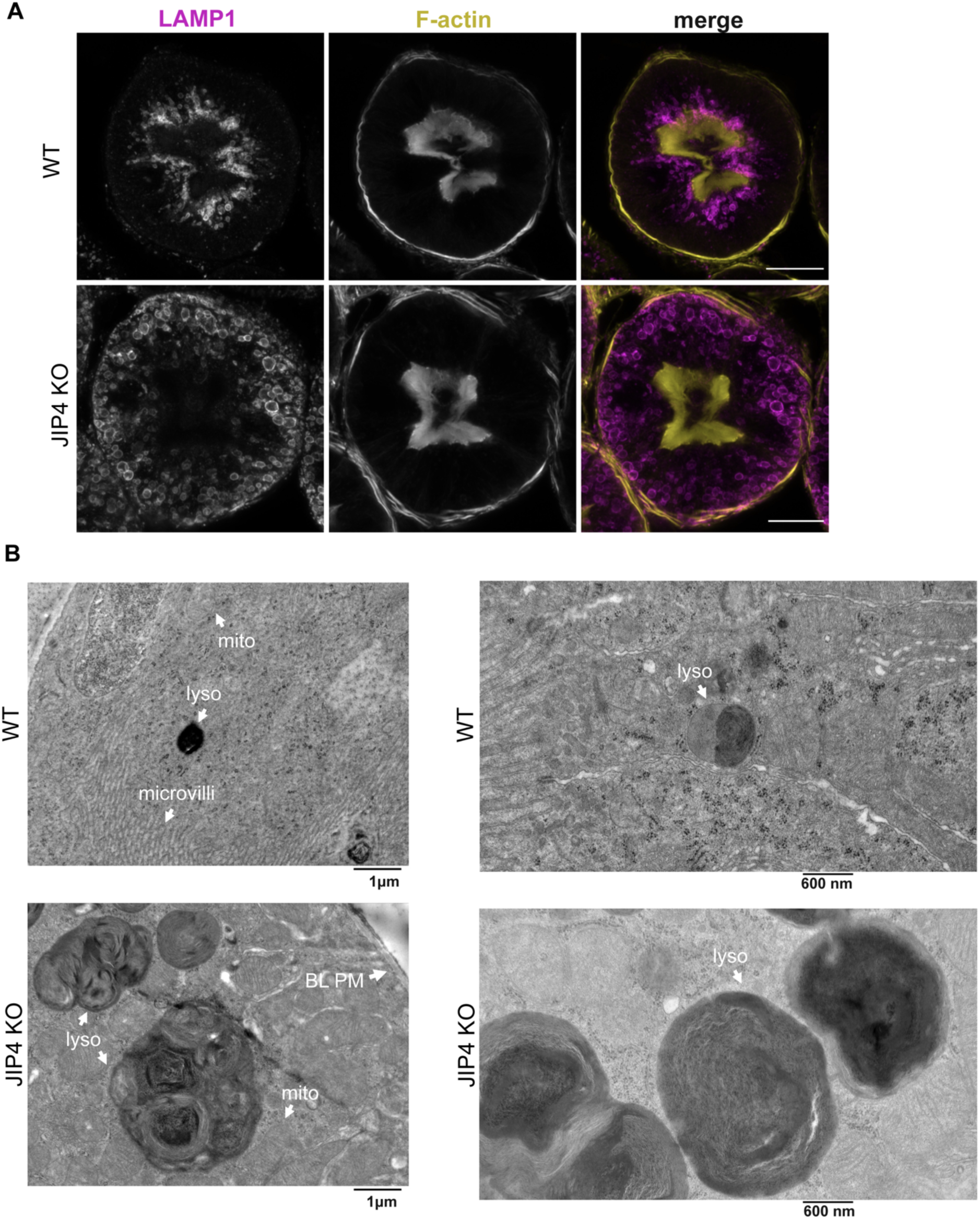
At 2 months, JIP4 KO proximal tubule lysosomes are mislocalized and multilamellar, related to Fig. 5. **a,** Immunohistochemistry of WT vs JIP4 KO renal proximal tubular cross-sections. Scale bar = 20 µm. **b,** Electron microscopy of WT vs JIP4 KO kidneys. Lyso = lysosome, mito = mitochondria, microvilli = apical plasma membrane microvilli, BL PM = basolateral plasma membrane.

## Supplemental Data

**Table S1:** Summary of antibodies used in this study.

| **Antibody** | **Acquired from** | **Reference number** | **Concentration** |
| --- | --- | --- | --- |
| FLAG | Cell Signaling Technologies | 2368,<br>RRID:AB_2217020) | WB: 1:5000<br>IF: 1:100 |
| JIP4 | Cell Signaling Technologies | 5519,<br>RRID:AB_10828724 | WB: 1:2500<br>IF: 1:100 |
| GAPDH | Encor Biotechnology | MCA-1D4,<br>RRID:AB_2107599 | WB: 1:60000 |
| HA | Cell Signaling Technologies | 3724S,<br>RRID:AB_1549585 | WB: 1:1000 |
| Vinculin | Cell Signaling Technologies | 13901,<br>RRID:AB_2728768 | WB: 1:10000 |
| LC3B | MBL International | PM036,<br>RRID:AB_2274121 | WB: 1:1000 |
| LAMP1 (H4A3) | DSHB | H4A3,<br>RRID:AB_2296838 | WB: 1:10000<br>IF: 1:1000 |
| TMEM55B | Proteintech | 23992-1-AP,<br>RRID:AB_2879391 | WB: 1:5000 |
| NEDD4 | Cell Signaling Technologies | 2740,<br>RRID:AB_2149312 | WB: 1:5000 |
| RFP | Rockland | 600-401-379,<br>RRID:AB_2209751 | WB: 1:10000 |
| LAMP1 (D2D11) | DSHB | 9091S,<br>RRID:AB_2687579) | WB: 1:10000<br>IF: 1:1000 |
| anti-megalin/LRP2 | Gift from Dr Daniel Biemsderfer and Dr Peter Aronson; Yale University | MC-220 ,<br>RRID:AB_3696948 | WB: 1:1000<br>IF: 1:100 |
| Cathepsin D | R&D Systems | MAB1029,<br>RRID:AB_2292411 | WB: 1:5000<br>IF: 1:100 |
| Clathrin Light Chain | EMD Millipore / Sigma | AB9884,<br>RRID:AB_11211734 | WB: 1:1000<br>IF: 1:100 |
| Rat IgG (HRP) | Cell Signaling Technologies | 7077S,<br>RRID:AB_10694715 | 1:2000 |
| Rabbit IgG (HRP) | Cell Signaling Technologies | 7074S,<br>RRID:AB_2099233 | 1:2000 |
| Mouse IgG (HRP) | Cell Signaling Technologies | 7076S,<br>RRID:AB_330924 | 1:2000 |
| AlexaFluor 488 anti-rat | Invitrogen | A21208,<br>RRID:AB_2535794 | 1:200 |
| Alexa 488-Rabbit | Invitrogen | A21206,<br>RRID:AB_2535792 | 1:200 |
| Alexa 568-Rabbit | Invitrogen | A10042,<br>RRID:AB_2534017 | 1:200 |
| Alexa 647-Mouse | Invitrogen | A21202,<br>RRID:AB_141607 | 1:200 |
| Alexa 594-Goat | Invitrogen | A11058,<br>RRID:AB_2534105 | 1:200 |

**Table S2:**
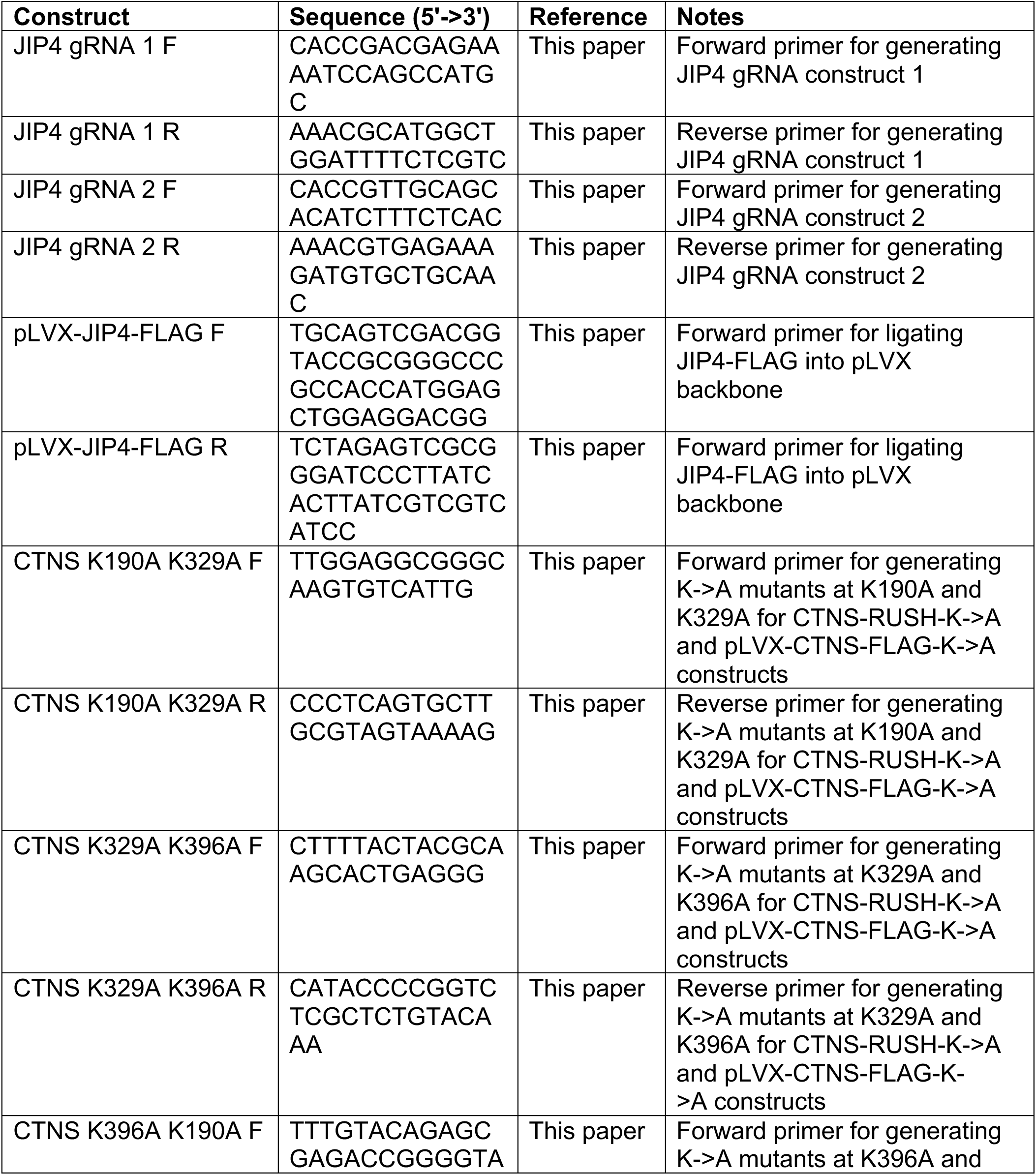
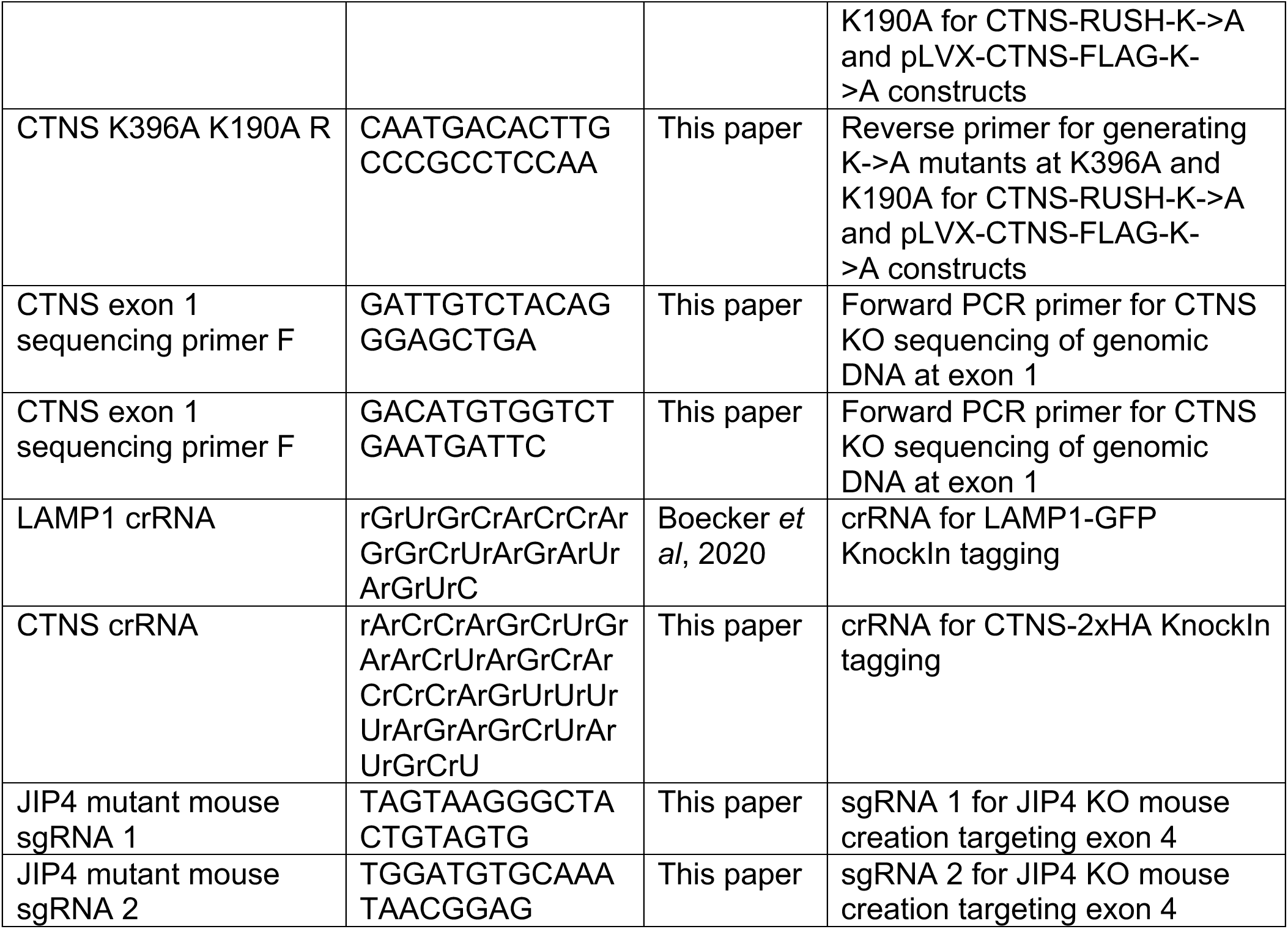
Summary of primers oligonucleotide sequences and crRNA used in this study.

**Table S3:**
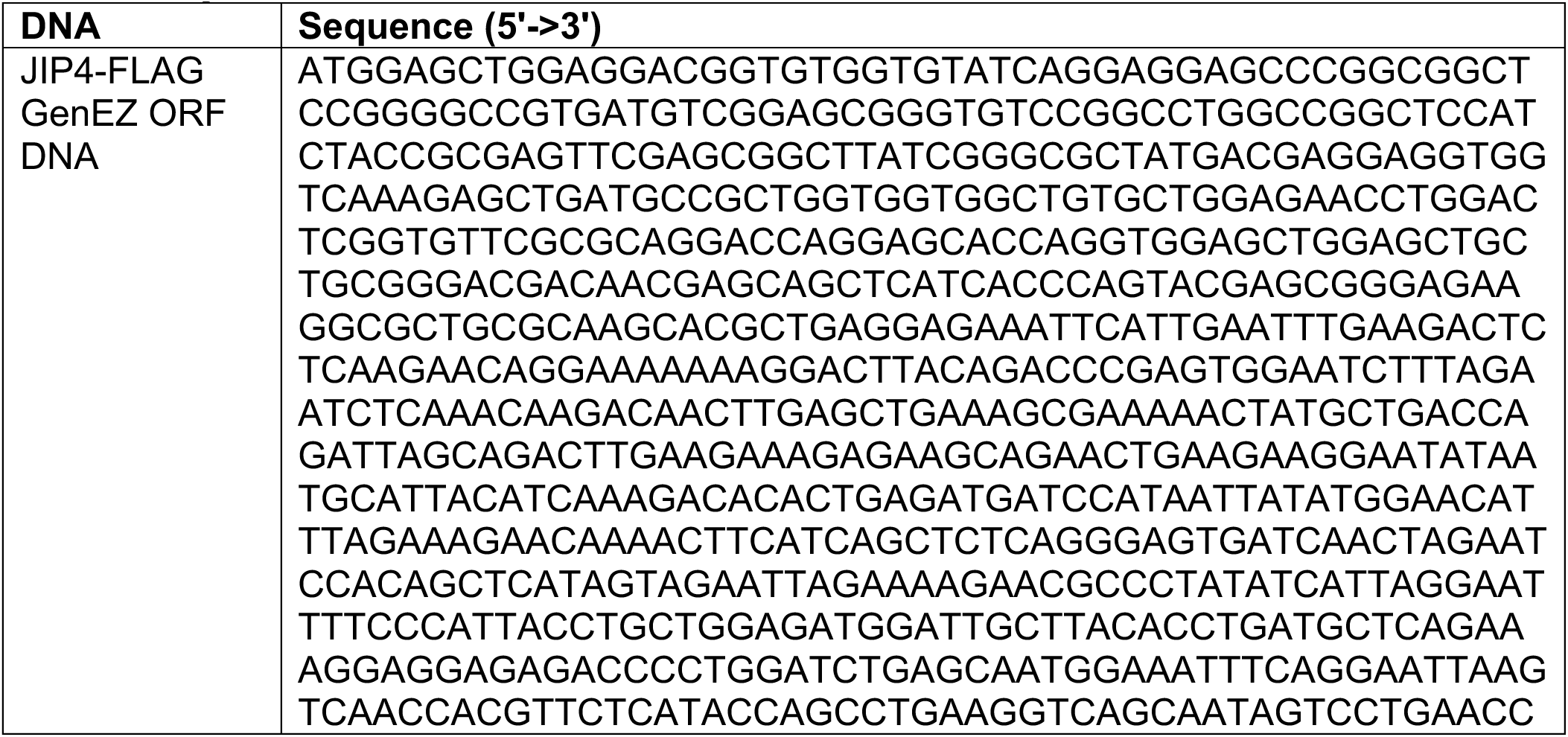
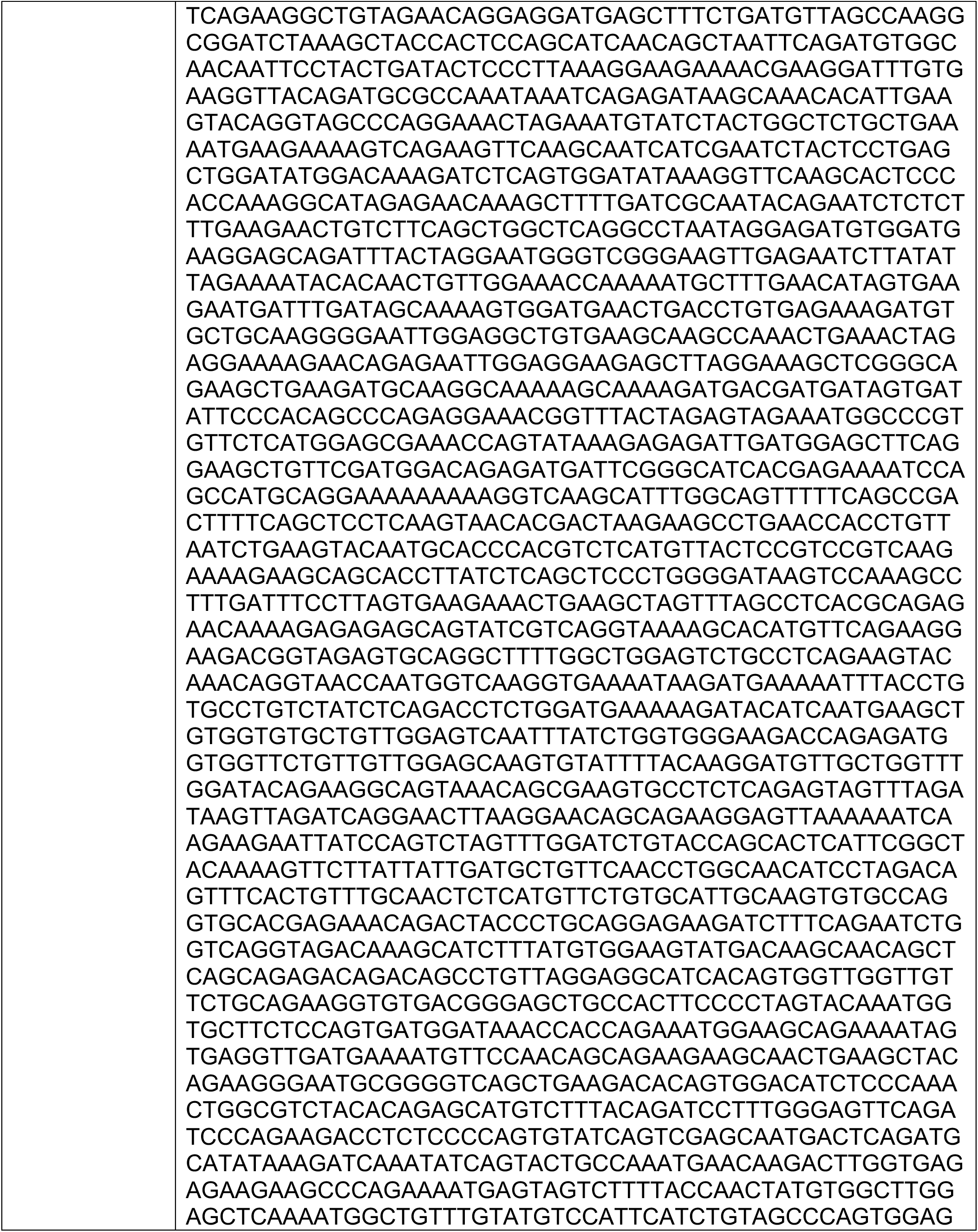
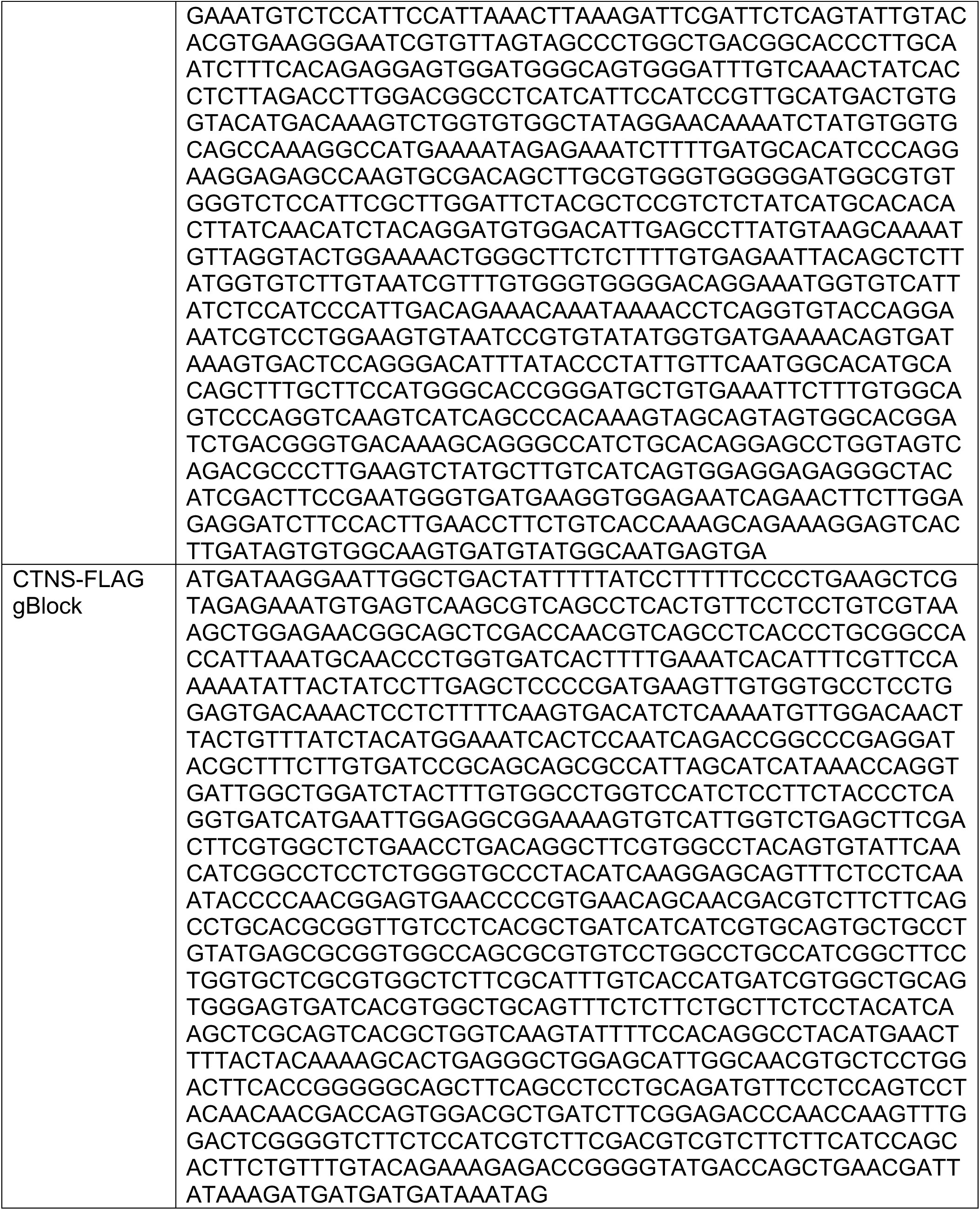
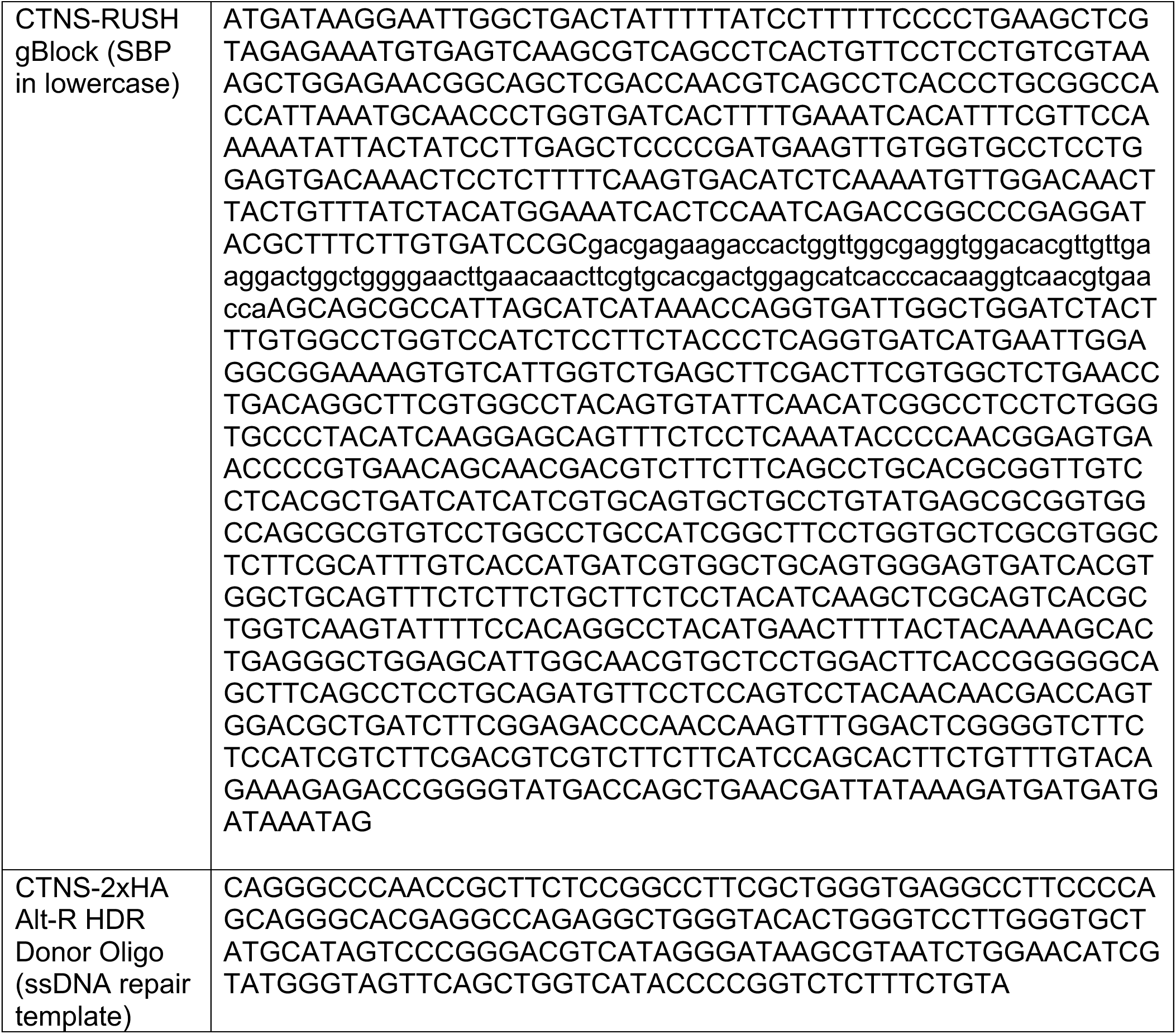
Summary of double stranded DNA blocks and ssDNA sequences used in this study.

**Table S4:** Summary of siRNAs used in this study.

| siRNA | Supplier | Reference |
| --- | --- | --- |
| Dharmacon ON-TARGETplus Non-Targeting siRNA Pool | Horizon Biosciences | D-001810-10-05 |
| Dharmacon OnTarget Plus siRNA smartPOOL TMEM55A | Horizon Biosciences | L-013808-01-0005 |
| Dharmacon OnTarget Plus siRNA smartPOOL TMEM55B | Horizon Biosciences | L-016425-02-0005 |

## Notes

### Competing Interest Statement

The authors have declared no competing interest.

